# Plant-mediated effects of fire and fragmentation drive plant-pollinator interaction β-diversity in fire-dependent pine savannas

**DOI:** 10.1101/2023.08.01.551484

**Authors:** Pablo Moreno-García, Johanna E. Freeman, Joshua W. Campbell, Eben N. Broadbent, Angélica M. Almeyda Zambrano, Gabriel Prata, Danilo R. A. de Almeida, Scott Gilb, Benjamin Baiser

## Abstract

Interaction β-diversity is an essential measure to understand and conserve species interactions and ecosystem functioning. Interaction β-diversity explains the variation in species interactions across spatial and temporal gradients, resulting from species turnover or interaction rewiring. Each component of interaction β-diversity has different ecological implications and practical consequences. While interaction β-diversity due to species turnover is related to assembly processes and fragmentation, rewiring can support high biodiversity and confer resilience to ecological networks. Despite this, it is unclear whether both components respond to the same or different ecological drivers. Here, we assessed the ecological drivers of plant-pollinator interaction β-diversity and its components across 24 sites in 9 Longleaf Pine (LLP) savannas in north and central Florida. We evaluated the effects of flowering plant composition and flower abundance, vegetation, fire regime, soil moisture, terrain characteristics, climate, spatial context, and geographic location. We used path analysis to evaluate the drivers of spatial interaction β-diversity and its main components. We then used generalized linear mixed models to assess the temporal patterns of spatial β-diversity among sites within preserves. We found that plant-pollinator networks in LLP savannas are highly variable across space and time, mainly due to species turnover and possibly in response to abiotic gradients and dispersal boundaries. Flower abundance and flowering plant composition, geographic location, fire seasonality, soil moisture, and landscape context were the main drivers of plant-pollinator β-diversity, highlighting the role of fire management and habitat connectivity in preserving plant-pollinator networks.

## INTRODUCTION

Variation in species composition across spatio-temporal scales, also known as β-diversity (Whittaker 1972; 1960), is a key component of regional species diversity and an important measure for assessing conservation efforts (Socolar et al. 2016). For example, β-diversity can be used to evaluate biotic homogenization (McKinney and Lockwood 1999; Li et al. 2020), community responses to anthropogenic pressures (Nascimbene and Spitale 2017; Santana et al. 2017), or prioritization of conservation areas (Zamora, Verdú, and Galante 2007; Ruhí, Datry, and Sabo 2017). Studies of β-diversity have commonly focused on species, but recent advances have extended the β-diversity framework to ecological interactions (Novotny 2009; Poisot et al. 2012; Fründ 2021). Species interactions differ from simple species co-occurrence even though co-occurrence is required and sometimes a good predictor of interactions (e.g., Harris 2016; e.g., Zurell, Pollock, and Thuiller 2018). The particular context of co-occurrence can govern the probability of an interaction (Poisot, Stouffer, and Gravel 2015; Burkle, Myers, and Travis Belote 2016). For example, species interactions vary with species densities (CaraDonna et al. 2017; de Manincor et al. 2020), phenology (CaraDonna et al. 2017; Morente-López et al. 2018; de Manincor et al. 2020), presence of other species (Traveset and Richardson 2006), and environmental characteristics (Burkle and Alarcón 2011; Poisot et al. 2017). Thus, species interaction β-diversity is composed of spatial or temporal differences in species co-occurrences (i.e., species turnover) and differences despite uniform species co-occurrences (i.e., interaction rewiring; Fig. 1) (Poisot et al. 2012).

**Figure 1.**
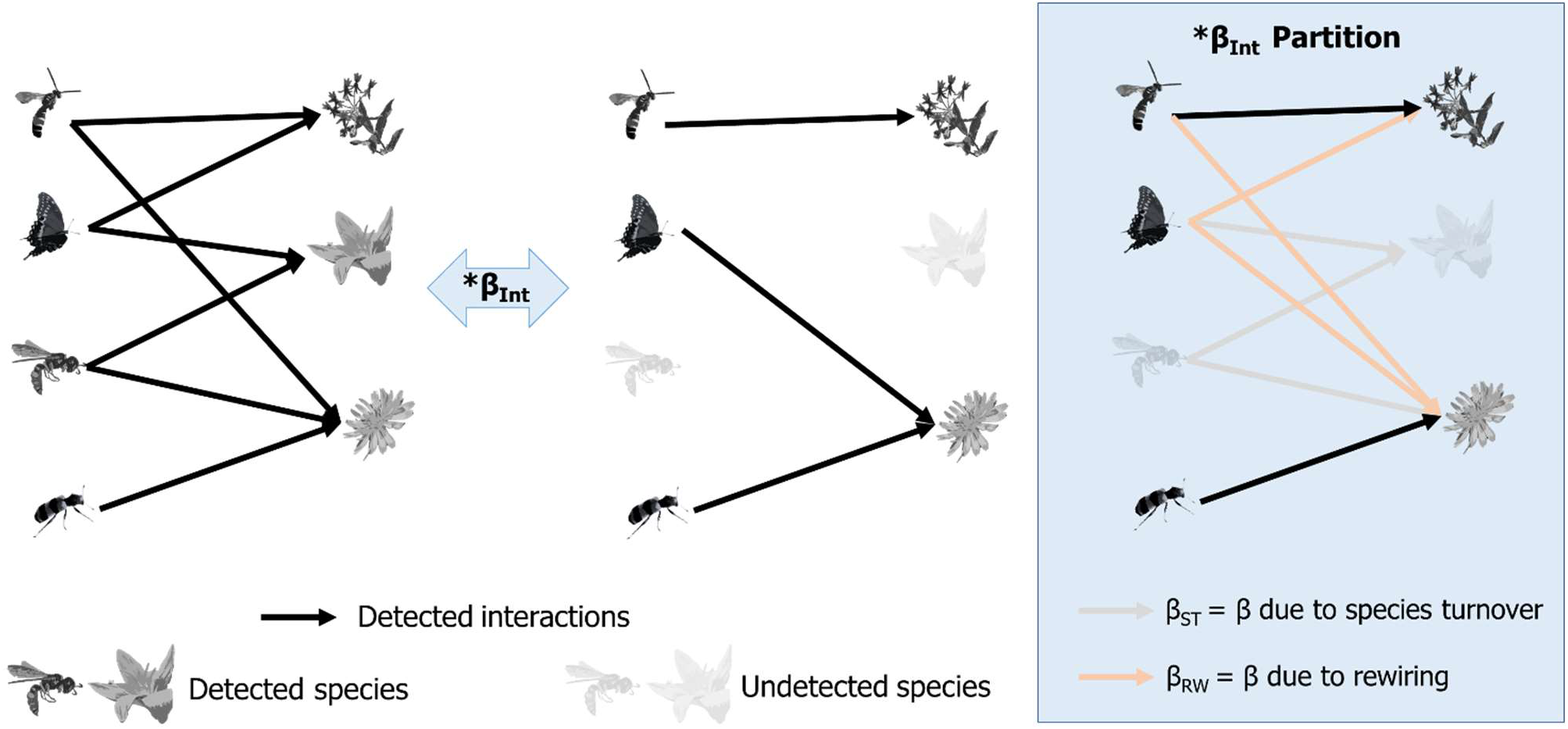
Graphic explanation of interaction β-diversity and its partition among the contributions of species turnover and rewiring. The two networks on the left are being compared, the blue box on the right details the contributions of species turnover and rewiring (it details the process characterized by the blue arrow). Detected species are highlighted, while missing or undetected species appear faded. Black lines represent the detected interactions, grey ones represent interaction variation due to species turnover, and orange ones represent interaction variation due to rewiring (variation despite species being detected in both samples).

The distinction between interaction β-diversity due to species turnover and interaction rewiring has practical implications for conservation (Burkle, Myers, and Travis Belote 2016; Fründ 2021). The relative contributions of species turnover and interaction rewiring are associated with species specialization (Kaiser-Bunbury et al. 2010; CaraDonna et al. 2017). Generalist species can participate in different interactions depending on the local context, resulting in partner switching and interaction rewiring. In addition to specialization, the participation of species in context-dependent interactions and subsequent increase in rewiring may depend on the cognitive, morphological, taxonomic, or demographic characteristics of a species (CaraDonna et al. 2017). The relative contributions of species turnover and interaction rewiring are also associated with specific causal mechanisms (CaraDonna et al. 2017) and ecological outcomes (Valdovinos et al. 2013). The contribution of species turnover tends to be higher during the assembly or disassembly of networks (CaraDonna et al. 2017; e.g., Kaiser-Bunbury et al. 2010) or following a disturbance (Burkle, Belote, and Myers 2022; e.g., Kaiser-Bunbury et al. 2010), while interaction rewiring confers network robustness to extinctions (Kaiser-Bunbury et al. 2010; Valdovinos et al. 2013) and promotes diversity by increasing resource partitioning and species coexistence (Valdovinos et al. 2013; Song and Feldman 2013). While the causal mechanisms and conservation consequences of interaction β-diversity due to species turnover and rewiring differ, it is unclear whether both components are driven by the same or different ecological variables. Few studies have assessed the ecological drivers of interaction β-diversity (Burkle, Belote, and Myers 2022), and they have found either consistent (e.g., CaraDonna et al. 2017) or inconsistent effects (Burkle, Belote, and Myers 2022) from the same ecological drivers. Understanding the drivers of each component is, however, important for the conservation of species interactions, especially in highly threatened ecosystems.

Fire-dependent longleaf pine savannas of the North American Coastal Plain (NACP), are highly diverse ecosystems that are now reduced to less than 5% of their original range due to fragmentation and fire suppression (Heuberger and Putz 2003; Van Lear et al. 2005; Platt 2007; Moylett, Youngsteadt, and Sorenson 2020). The NACP has recently been recognized as one of the Earth’s biodiversity hotspots, and much of the species diversity and endemism that support this designation are contributed by the understories of fire-dependent longleaf pine savannas (Noss et al. 2015). The plant-communities of frequently-burned pine savannas, which are very rich in insect-pollinated herb and shrub species, exhibit high, turnover-dominated β-diversity patterns (Freeman et al. 2019). Large-scale conservation of fragmented preserve networks in this region stands to benefit from a plant metacommunity approach that incorporates β-diversity as a central guiding concept, since different preserves harbor different components of the regional plant metacommunity (Freeman et al. 2019). Given the high β-diversity known to exist among flowering plant communities in this region, we would expect to see similarly high β-diversity in pollinator communities and their mutualistic interactions with flowering plants. The little research that has been conducted on plant-pollinator networks in longleaf pine savannas has shown that landscape-level factors (i.e., habitat fragmentation) play a large role in network architecture (Spiesman and Inouye 2013). Developing a better understanding of the patterns and drivers of plant-pollinator interaction β-diversity in NACP savannas at multiple scales will support conservation strategies at both the regional and local level.

In order to better understand mutualistic plant-pollinator interaction networks in the longleaf pine savanna region, we studied plant-pollinator interactions across a large regional sample of 9 preserves in north-central Florida throughout an entire flowering season; collected a suite of variables related to land management, abiotic conditions, and floral resources; and evaluated the causal pathways of interaction β-diversity and its partitions (i.e., interaction β-diversity due to species turnover and rewiring). Our overarching objective was to determine the roles of biotic and abiotic variables in driving plant-pollinator interaction β-diversity across space and time, including fire regime (frequency and seasonality of prescribed fires), landscape connectivity, savanna vegetation structure, soil moisture, geographic location, flower species composition, and flower abundance. We tested three hypotheses regarding the causal pathways between these variables: A) Direct effects (each variable directly affects the β-diversity of interactions); B) Flower resources mediation (flower abundance is the direct driver of interaction β-diversity, while the other variables are indirect drivers via their effects on flower abundance); and C) Serial vegetation structure-flower resources mediation (flower abundance is the direct driver of interaction β-diversity, vegetation structure is the direct driver of flower abundance, and all other variables are indirect drivers via their effects on vegetation structure) (Fig. 2).

**Figure 2.**
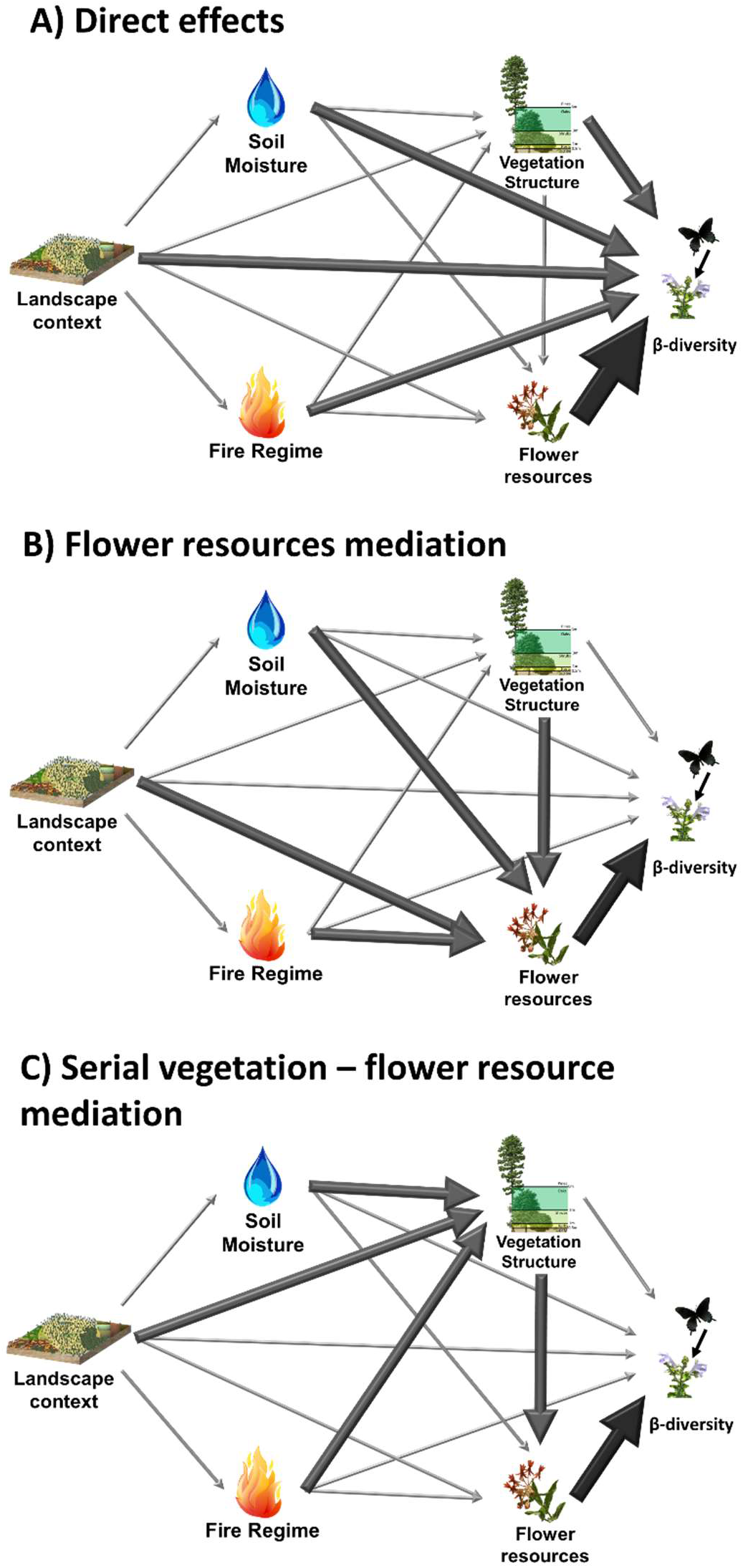
Schematic causal graph of relationships between environmental distances and β-diversity. Both arrow color and width emphasize the link strength. Β-diversity may refer to plant-pollinator interactions or its components. Flower resources include flower abundance and flowering plant composition dissimilarities. Vegetation includes vegetation LAI and structure distances. Fire management includes dissimilarities based on the number of prescribed fires and fire seasonality across the growing and dormant seasons. Soil moisture refers to differences in soil moisture. Landscape context includes weather, surrounding land uses, terrain, and geographic distances. The panels indicate the possible pathways by which landscape context, fire regime, and soil moisture may affect plant-pollinator interaction β-diversity via vegetation structure and flower resources.

## METHODS

### Study system

Pine savannas are fire-dependent ecosystems located in the North American Coastal Plain, which are characterized by a bi-layered vegetation with an open pine-oak canopy and a highly diverse herbaceous layer (Myers 1990; Reinhart and Menges 2004; Collins et al. 2006; Platt 2007; Menges 2007; Carr, Robertson, and Peet 2010). Pine savannas are maintained by frequent fires (i.e., fire return intervals of 1-5 years) (Heuberger and Putz 2003; Menges 2007; Freeman et al. 2017) and in the absence of fire, hardwood encroachment leads to litter accumulation, reduced light penetration, and the formation of closed-canopy oak forests with depauperate herbaceous communities (Menges and Hawkes 1998; Platt 2007; Hiers et al. 2007; Carr et al. 2009; Britez, Romero, and Putz 2020). While mesic savannas tend to foster more diverse plant communities (Myers 1990; Freeman et al. 2017), they are more susceptible to oak encroachment (Freeman et al. 2019). Fragmented savannas are also more prone to fire suppression and subsequent degradation (Heuberger and Putz 2003), in a process fueled by past and prospective human development in the area (Heuberger and Putz 2003; Boda 2018). In north and central Florida pine savannas, flowers and pollinators tend to be present and active for the entire growing season, from March to October. Flowering plants in pine savannas display two bloom peaks occurring in spring (April-May) and fall (August-October) (Platt, Evans, and Davis 1988). Pollinators tend to be especially active during the fall flowering peak, when most composite species bloom (Asteraceae, Magnoliopsida: Euasterids) (Platt, Evans, and Davis 1988; Hall and Ascher 2011; Atwater 2013).

### Study sites

We studied nine pine savannas (i.e., preserves) across three different regions in north and central Florida (Supplementary Material 1). All savannas are located within historical Timucua and Seminole native territories, additionally two of the preserves are located within historical Tocobaga native territory (Temprano 2015). Early records indicate that these sites were frequently burned by wildfires and Native Peoples prior to European colonization (Davison and Bratton 1987; Van Lear et al. 2005). We established either two or three 100 m x 100 m survey sites within each preserve in areas that had been last burned between October 2017 and July 2018. Study sites were a minimum of 1 km apart, following Spiesman and Inouye (2013). Within each one-hectare site, we established five 10 m x 10 m flower abundance sampling plots, two flower abundance sampling transects bisecting the plot N-S and E-W, and 25 5 m x 5 m vegetation quads.

### Pollination surveys

#### Sampling

We carried out monthly plant-visitor surveys on each of our sites from March to October 2019. We divided each 100 m x 100 m research site into four quadrants and surveyed each quadrant for 30 minutes, following semi-meandering variable transects (VT, Nielsen et al. 2011). Observers captured insect visitors that contacted any flower reproductive structures along the transects and recorded the location, plant species, and time of capture. We collected the insect samples for their posterior identification ex situ; thus, our data portrays interaction frequency rather than strength (i.e., we only include one visit per pollinator individual). We limited our observations to optimal times and climatic conditions (sensu Abreu and Vieira 2004; Bartomeus, Bosch, and Vilà 2008). We sampled one preserve per day, rotating the site order monthly.

#### Sample treatment and identification

We then identified the samples to the lowest possible taxonomic level, using microscopy, and the taxonomic guides detailed on Supplementary Material 2. We used Leica models MZ16 (10x lens) and M80 (1.0x Achromat) microscopes, with total magnification of 115x and 60x, respectively. We were assisted by taxonomic experts from the Florida State Collection of Arthropods, US Department of Agriculture, and the University of Florida.

### Measurements of β-diversity

We evaluated interaction β-diversity following the conceptual framework in Poisot et al. (2012) and methods in Fründ (2021) (i.e., betalinkr in bipartite v. 2.18 in r v. 4.0.3, Dormann 2020). We compiled monthly and annual by-site and by-preserve plant-pollinator interactions in incidence matrices, in which each element *i_ij_* indicates the number of pollinators of species *j* found on plant *i* (Chacoff, Resasco, and Vázquez 2018). Then, we used Ružička dissimilarities to evaluate β-diversity of interactions (β^Int^). Ružička is equivalent to Jaccard dissimilarities for weighted data, in this case number of interactions. We partitioned interaction β-diversity (β_Int_) into rewiring (β_RW_) and interaction β-diversity due to species turnover (β_ST_) (Fig. 1). We calculated measures of interaction β-diversity and its components (i.e., β_Int_, β_ST_, and β_RW_) among our plant-pollinator networks at two spatial (i.e., site, preserve) and temporal scales (i.e., monthly, and annual).

### Independent variables

We evaluated the relationship between interaction β-diversity and asset of independent ecological predictors. We used ecological distances to explain spatial patterns of β_Int_, β_ST_, and β_RW_ using path analysis. In the path analysis, we assessed the effects of flowering plant abundance and diversity, leaf area index (LAI) across four vegetation strata, land cover and vegetation structure, fire regime (i.e., prescribed fire frequency and seasonal diversity), soil moisture, weather, terrain, landscape, and geographical context (Table 1) on interaction β-diversity. We calculated all ecological distances using the package vegan v. 2.6.4 in R v.4.0.3 (Dixon 2003). Furthermore, we used monthly flowering plant resources to evaluate the monthly variation of spatial interaction β-diversity.

**Table 1.**
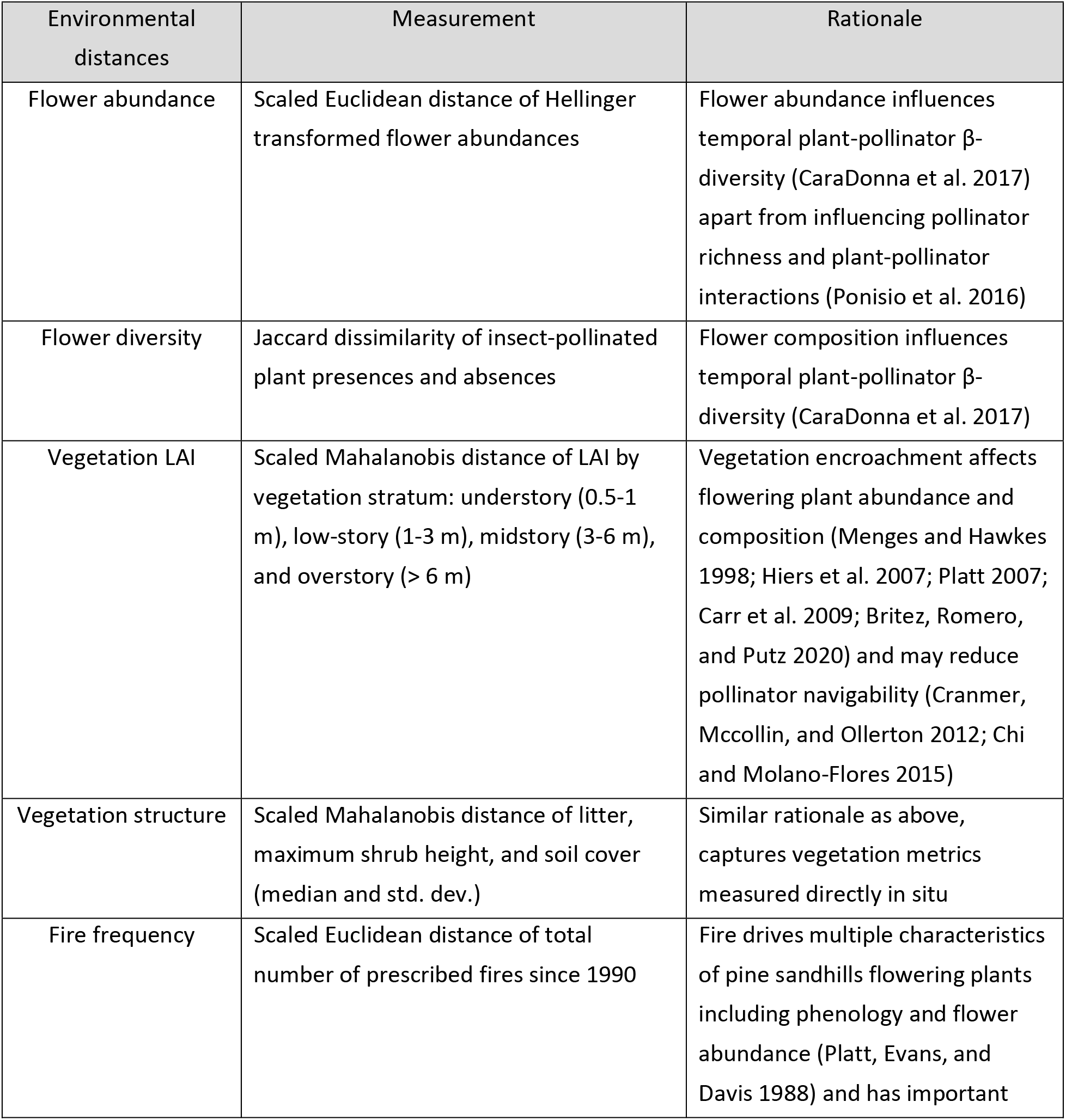
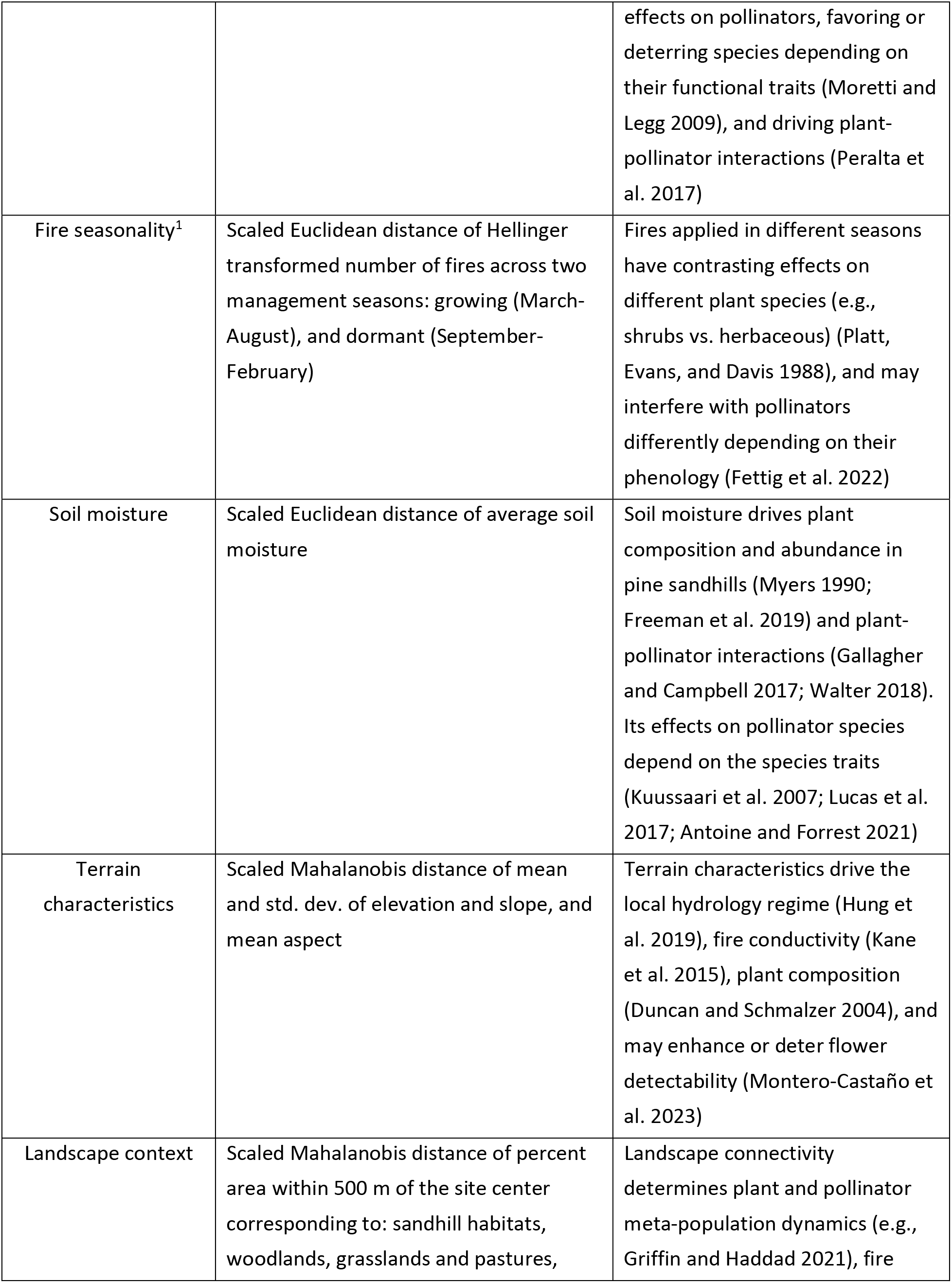
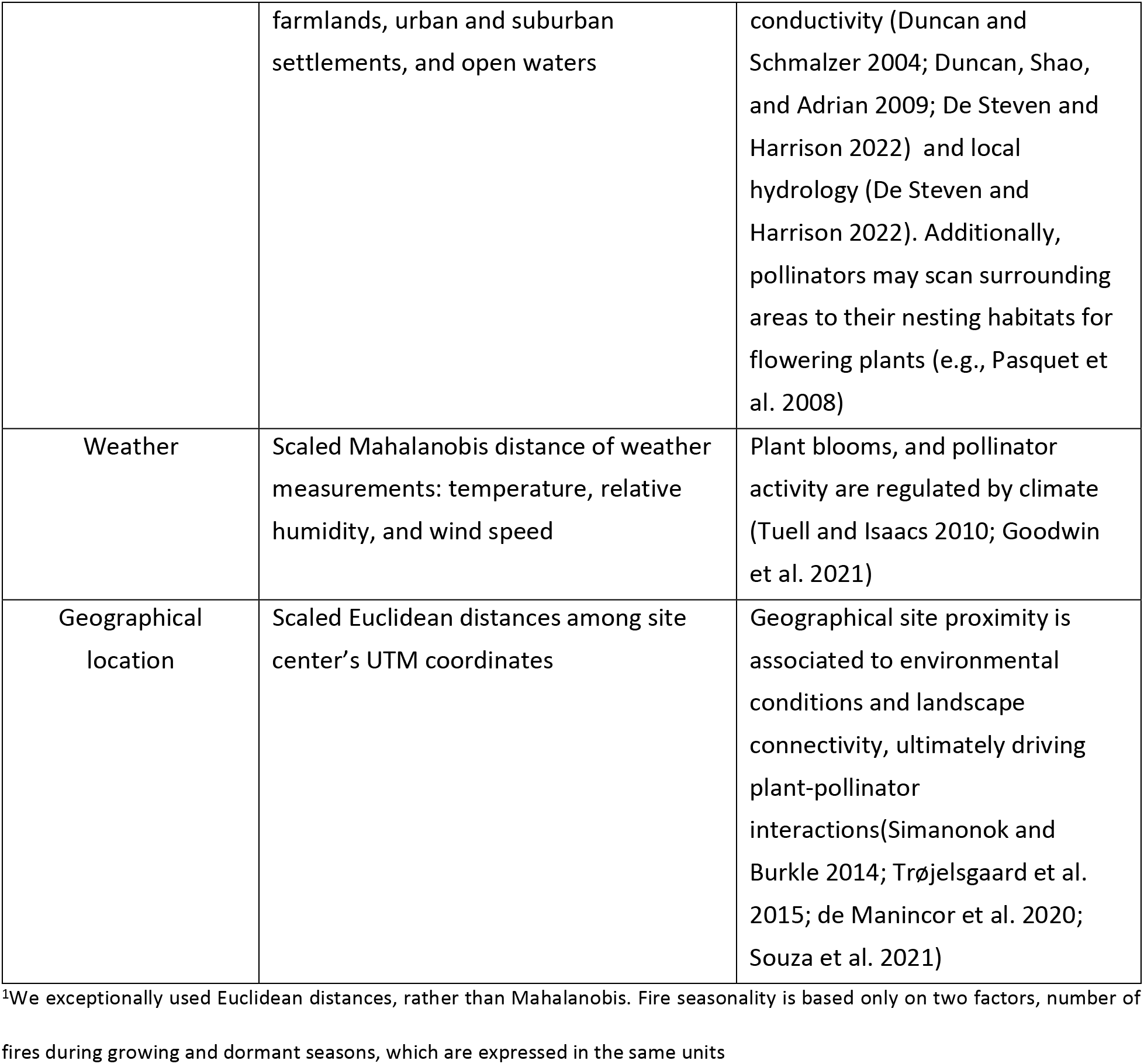
Description of the selected ecological distances. We registered monthly observations of flower abundance (and flowering plant diversity), soil moisture, and weather characteristics, and pooled them for further analyses. We assessed vegetation LAI, terrain characteristics, and landscape context via aerial LiDAR surveys. We derived fire frequency and seasonality from the history of prescribed fires and mechanical treatments that we obtained from preserve managers. We gathered site UTM coordinates, and vegetation diversity and structure data in situ.

| Environmental distances | Measurement | Rationale |
| --- | --- | --- |
| Flower abundance | Scaled Euclidean distance of Hellinger transformed flower abundances | Flower abundance influences temporal plant-pollinator β-diversity (CaraDonna et al. 2017) apart from influencing pollinator richness and plant-pollinator interactions (Ponisio et al. 2016) |
| Flower diversity | Jaccard dissimilarity of insect-pollinated plant presences and absences | Flower composition influences temporal plant-pollinator β-diversity (CaraDonna et al. 2017) |
| Vegetation LAI | Scaled Mahalanobis distance of LAI by vegetation stratum: understory (0.5-1 m), low-story (1-3 m), midstory (3-6 m), and overstory (> 6 m) | Vegetation encroachment affects flowering plant abundance and composition (Menges and Hawkes 1998; Hiers et al. 2007; Platt 2007; Carr et al. 2009; Britez, Romero, and Putz 2020) and may reduce pollinator navigability (Cranmer, Mccollin, and Ollerton 2012; Chi and Molano-Flores 2015) |
| Vegetation structure | Scaled Mahalanobis distance of litter, maximum shrub height, and soil cover (median and std. dev.) | Similar rationale as above, captures vegetation metrics measured directly in situ |
| Fire frequency | Scaled Euclidean distance of total number of prescribed fires since 1990 | Fire drives multiple characteristics of pine sandhills flowering plants including phenology and flower abundance (Platt, Evans, and Davis 1988) and has important |
|  |  | effects on pollinators, favoring or deterring species depending on their functional traits (Moretti and Legg 2009), and driving plant-pollinator interactions (Peralta et al. 2017) |
| Fire seasonality<sup>1</sup> | Scaled Euclidean distance of Hellinger transformed number of fires across two management seasons: growing (March-August), and dormant (September-February) | Fires applied in different seasons have contrasting effects on different plant species (e.g., shrubs vs. herbaceous) (Platt, Evans, and Davis 1988), and may interfere with pollinators differently depending on their phenology (Fettig et al. 2022) |
| Soil moisture | Scaled Euclidean distance of average soil moisture | Soil moisture drives plant composition and abundance in pine sandhills (Myers 1990; Freeman et al. 2019) and plant-pollinator interactions (Gallagher and Campbell 2017; Walter 2018). Its effects on pollinator species depend on the species traits (Kuussaari et al. 2007; Lucas et al. 2017; Antoine and Forrest 2021) |
| Terrain characteristics | Scaled Mahalanobis distance of mean and std. dev. of elevation and slope, and mean aspect | Terrain characteristics drive the local hydrology regime (Hung et al. 2019), fire conductivity (Kane et al. 2015), plant composition (Duncan and Schmalzer 2004), and may enhance or deter flower detectability (Montero-Castaño et al. 2023) |
| Landscape context | Scaled Mahalanobis distance of percent area within 500 m of the site center corresponding to: sandhill habitats, woodlands, grasslands and pastures, | Landscape connectivity determines plant and pollinator meta-population dynamics (e.g., Griffin and Haddad 2021), fire |
|  | farmlands, urban and suburban settlements, and open waters | conductivity (Duncan and Schmalzer 2004; Duncan, Shao, and Adrian 2009; De Steven and Harrison 2022) and local hydrology (De Steven and Harrison 2022). Additionally, pollinators may scan surrounding areas to their nesting habitats for flowering plants (e.g., Pasquet et al. 2008) |
| Weather | Scaled Mahalanobis distance of weather measurements: temperature, relative humidity, and wind speed | Plant blooms, and pollinator activity are regulated by climate (Tuell and Isaacs 2010; Goodwin et al. 2021) |
| Geographical location | Scaled Euclidean distances among site center's UTM coordinates | Geographical site proximity is associated to environmental conditions and landscape connectivity, ultimately driving plant-pollinator interactions (Simanonok and Burkle 2014; Trøjelsgaard et al. 2015; de Manincor et al. 2020; Souza et al. 2021) |

<sup>1</sup>We exceptionally used Euclidean distances, rather than Mahalanobis. Fire seasonality is based only on two factors, number of fires during growing and dormant seasons, which are expressed in the same units

#### Flowering plant abundance and diversity distances

We derived dissimilarity matrices from flower abundance, and flowering plant diversity surveys. We conducted two different monthly surveys of flowering plants in each of our sites from March to October 2019, alongside plant-pollinator surveys. In one survey, we censused flowering plants in the five 10 m x 10 m sampling plots. In the second, we censused flowering plants along the N-S and W-E transects. The transects spanned 45 m from the center to the edge of the site at each main cardinal point, excluding the overlapping area with the central sampling plot (5 m in each transect). We surveyed the plants within a meter to each side of the transect; thus, we surveyed an area of 45 m x 2 m for each transect (i.e., four transects from the center to each cardinal point). We used the same sampling method for both surveys, counting and summing all the stems with blooming flowers. Hereafter, when we mention flower abundance, we refer to the abundance of blooming flowering plants, so as to avoid the confusion between the abundance of blooming and overall present angiosperms (i.e., flowering plants). We summed monthly flower abundances by site, to obtain annual flower abundances. We calculated the scaled Euclidean distance of the Hellinger-transformed flower abundances among study sites.

The vegetation at each research site was very heterogeneous and patchy. To obtain a truly representative plant community sample, we surveyed the full plant community in a grid of 25 5 m x 5 m quads laid out evenly across the hectare. The full plant survey was conducted between September and November 2019. All plants in the 5 m x 5 m quad were identified to the lowest taxonomic level possible and presence/absence was recorded for each species. Then, we sampled insect-pollinated plants presence within each of the vegetation quads. We aggregated the plant species samples from the 25 individual quads into a single presence/absence dataset for each research site. We added the plant species documented in the monthly flower surveys, but not in the final vegetation sample (i.e., ephemerals). We computed the Jaccard distance of the flowering plant presences to evaluate differences in species diversity.

#### Vegetation LAI and structure distances

We collected dense LiDAR data over each site using the drone-borne GatorEye Unoccupied Flying Laboratory (www.gatoreye.org). The GatorEye LiDAR system incorporates a Phoenix Scout ultra-core with a Velodyne VLP-32 sensor. We collected LiDAR data at a speed of 10 m/s and a height of 100 m aboveground level. We spaced the flightlines 20 meters apart and processed the data against a base station, providing us with an absolute geolocation precision of less than 5 cm. Then, we cleaned and processed the LiDAR point clouds to standard deliverables of Digital Elevation Model, Digital Surface Model, and Canopy Height Model raster images using the GatorEye Multi-Scalar Post-Processing (GMSPP v218). The output point clouds had a density of more than 500 pts/m^2^. We post-processed the LiDAR point clouds to obtain leaf area indices per strata, following the ‘GatorEye Advanced Post-Processing Workflow 1’ (GAPP-W1). We assessed the distribution of leaf area across key vertical height strata, specifically the grass/forb (0.5-1 m), shrub (1-3 m), subdominant tree (3-6 m), and dominant tree strata (> 6 m) (de Almeida et al. 2019; 2020). We did not estimate the LAI of plants shorter than 0.5 m to avoid any issues arising from the inaccurate estimation of digital elevation models for each site. We transformed absolute leaf area to leaf area indices (LAI) by dividing total leaf area by 1 ha. A number of proportions were greater than one, due to the overlap of vegetation at different heights. We used scaled Mahalanobis distance to assess pairwise differences in vegetation LAI across the four strata between study sites. Detailed system specifications and workflow descriptions are provided in the GatorEye overview manuscript (Broadbent et al. 2021 v22, downloaded from www.gatoreye.org on 10/26/2021).

As another approach to quantify vegetative structure, we measured maximum shrub height, soil cover (percentage of shrubs, herbs, litter, and bareground to the nearest 5%), litter height, and woody debris in each of our 25 5 x 5 m vegetation sampling plots. We assessed soil cover in the southwest quadrant, and litter height at the center of each vegetation sampling plot. Observers calibrated their estimates of soil cover percentages before starting the surveys. We calculated the site median and standard deviation of shrub and litter height, soil cover, and woody debris across the 25 vegetation quads in each site. We used the median values instead of the means to limit the influence of extreme measurements. We assessed collinearity and excluded median litter height and cover from the final analysis. We calculated the scaled Mahalanobis distance of the remaining variables.

#### Fire regime distances

To construct 30-year fire and management histories for each study site (including the dates and details of all known fires and fire surrogate treatments), we distributed a detailed questionnaire to the managing agencies of each site. The total number of fires per site ranged from 4 to 15, with a mean of 7.6 fires since 1990. A small number of fire surrogates (e.g., mechanical thinning, hack-and-squirt herbicide) were also used to reduce hardwood encroachment on some sites (a total of 10 surrogate treatments vs. 183 fires over the whole study). Although fire surrogates differ from fires in some important ways, this type of fire history metric best describes the understory when fire surrogate treatments are simply counted as fires (Freeman et al. 2019). Hereafter, when we refer to fires and the fire regime, we are also referencing the small number of fire surrogates and including those fire surrogate treatments as part of the fire regime calculations. We evaluated the number of fires, as well as fire frequency across the most commonly used management seasons: growing (March – August) and dormant season (September – February). We used scaled Euclidean distances for the number of fires and fire seasonality. We used Hellinger transformed fire numbers by season to evaluate fire seasonality (vegan 2.6-4 in R v. 4.0.3, Dixon 2003).

#### Soil and terrain distances

We conducted monthly surveys of soil moisture using a FieldScout TDR 350 Soil Moisture Meter with 7.6 cm rods. During each survey, we took five measurements of soil moisture in the five 10 m x 10 m sampling plots. We averaged the measurements (5 measurements x 5 plots) to obtain the monthly mean values per site. We averaged the monthly measurements by site to obtain annual means and used scaled Euclidean distances to evaluate pairwise differences of soil moisture between study sites. We assessed terrain characteristics in our aerial surveys, including elevation, slope, and aspect. We calculated the mean elevation, slope, and aspect, as well as the standard deviation of elevation and slope. We calculated the scaled Mahalanobis distance of terrain characteristics.

#### Landscape context distance

To assess the land use (LU) surrounding our sites, we delimited land cover polygons within a 500 m radius of each site center using high-resolution satellite imagery from Google Earth (years 2019-2021). We considered six different LU: pine savannas, woodland habitats, open grasslands and pastures, farmlands, urban and suburban areas, and open waters. We evaluated the proportion of each land use within 500 m of our site centers. We validated our imagery interpretation through field visits. Our sites were mostly surrounded by closed canopy natural forests (e.g., oak hammocks) and open canopy natural savanna. We calculated the scaled Mahalanobis distance of LU proportion within the surrounding landscape.

#### Weather distance

We carried out monthly weather surveys at each of the sites using a Kestrel 3500 Pocket Weather Meter. We measured temperature, relative humidity, and average windspeed *in situ,* at the center of each site. Then, we averaged the monthly values by site to obtain annual weather estimates. We used the scaled Mahalanobis distance of weather conditions.

#### Geographical distance

We recorded the UTM coordinates of our site centers within UTM zone 17N. We calculated the scaled Euclidean distances among our site centers (packages adespatial v.0.3.21, Dray et al. 2018; and vegan v.2.6.4 in r v.4.0.3, Dixon 2003).

#### Monthly flowering plant resources

We evaluated the effect of monthly flower abundance, and flowering plant diversity, and evenness on monthly spatial patterns of interaction β-diversity and its components within preserves (i.e., within-preserve among-site variation of interactions). We used the data on monthly flower abundance per site to characterize the monthly flower plant resources of each preserve utilizing two metrics: Chao 1 index of diversity (Chao 1987) and observed species richness. We calculated Chao 1 index of diversity using the monthly flower abundances per site and preserve (Chao 1987; rareNMtests v.1.2 in R v. 4.0.3, Cayuela and Gotelli 2014). We evaluated monthly flowering plant evenness using Pielou’s evenness J (Pielou 1969; vegan in v.2.6.4 R v.4.0.3, Dixon 2003).

### Statistical analyses

#### Path analysis: biogeographical and ecological drivers

We conducted path analyses to evaluate the direct and indirect effects of biogeographical and environmental dissimilarities on annual interaction β-diversity and its partition using permutation-based d-sep tests and permutation-based path analysis (Fourtune et al. 2018). These tests allowed us to evaluate whether sites that have more distinct environmental characteristics (e.g., soil moisture) are more likely to differ in their interactions (or its components), and whether this relationship is mediated by other environmental factors (e.g., flower abundance). Permutation-based d-sep tests are used to evaluate the overall fit of the data to the proposed causal graph, where p-values > 0.95 indicate a significant fit (if α=0.05). Permutation-based path analysis is used to evaluate the significance of each proposed path on the causal graph, where p-values < 0.05 indicate significance. We tested whether β_Int_, β_ST_, and β_RW_ depend on flower characteristics, vegetation structure, fire management, soil moisture, topography, landscape context, climate, and/or geography (Table 1, Fig. 2). We tested our hypothesized paths using multiple regression on distance matrices (MRM) (Fourtune et al. 2018). We compared 3 hypotheses (i.e., direct effects, flower resources mediation, and vegetation structure-flower resources mediation, Fig. 2) starting with the full model (all paths in Fig. 2) and selected the path model with the lowest AIC or the most parsimonious model with ΔAIC <2 from the model with the lowest AIC. Since network-derived metrics can be sensitive to sampling coverage (i.e., the probability of occurrence of the interactions), we repeated the analyses including only the sites with coverage > 0.6, balancing higher coverage levels with a sample of > 50% of the original sites (Supplementary Material 5). While a standard of 0.6 may not be ideal, coverages of 0.6 are not rare in plant-pollinator networks (e.g., Zackenberg, Olesen et al. 2008; Jordano 2016).

#### Monthly trends of spatial β-diversity

Plant-pollinator interactions vary along temporal gradients, such that spatial patterns of β-diversity (i.e., sites within a preserve) may not be consistent through time. We used generalized linear mixed models (GLMM) to evaluate whether the mean spatial variability of interaction β-diversity and its components within preserves varied predictably with monthly floral resources. In these models, either monthly β_Int_, β_ST_, and β_RW_ between sites within preserves served as response variables and monthly measures of flower abundances, Chao 1 diversity index, species richness, and Pielou’s J evenness index were predictor variables. We evaluated the random effects of region, preserve, and month on the intercept, as well as their additive and multiplicative effects following Zuur et al. (2009). The best-fit models did not include any random effects. Then, we selected the fixed structure by using backward model selection based on AIC values. We evaluated all parametric assumptions, including outliers, influential points, normality of residuals, heteroskedasticity, and linearity (car v.3.1.1 in R v. 4.0.3, Fox et al. 2016). We repeated the models with and without outliers to check for any inconsistencies. In the results we present the models without outliers (but see Supplementary Material 6 for the models with outliers). We repeated the analysis only including the preserves by month with coverage > 0.6 (Supplementary Material 5).

## RESULTS

Interaction β-diversity (β_Int_) was high across spatial and temporal units (i.e., surveys: 0.99 ± 0.02, sites: 0.97 ± 0.03, preserves: 0.94 ± 0.03, regions: 0.89 ± 0.01, and months: 0.98 ± 0.03), indicating that interactions are spatially and temporally structured in pine savannas. Interaction β-diversity due to species turnover (β_ST_) was very high among surveys (0.98 ± 0.07), sites (0.75 ± 0.14), and months (0.82 ± 0.21) where it explained most of β_Int_ but was lower among preserves (0.56 ± 0.14) and regions (0.35 ± 0.03). Interaction β-diversity due to rewiring (β_RW_) explained a large amount of β_Int_ among preserves and, especially, regions, where it surpassed β_ST_ (see Supplementary Material 6 for more information).

### Path analysis: biogeographical and ecological drivers

#### Plant-pollinator interaction β-diversity (β_Int_)

The best-fit SEM for β_Int_ (C-statistic = 38.455, df = 70, P-value = 0.999) described 51% of the variation in plant-pollinator interaction β-diversity (Fig. 3A) (Supplementary Material 3 Table 1) and supported the hypothesis of flower resources mediated effects (Fig. 2B). β_Int_ increased with differences in flower abundance (std. coef. = 0.454, P-value < 0.001) and flowering plant composition (std. coef. = 0.220, P-value = 0.001), as well as geographical distance (std. coef. = 0.162, P-value = 0.013). Hence, sites further away from one another and/or sites with different flower abundance and flowering plant composition tend to have different plant-pollinator interactions.

**Figure 3.**
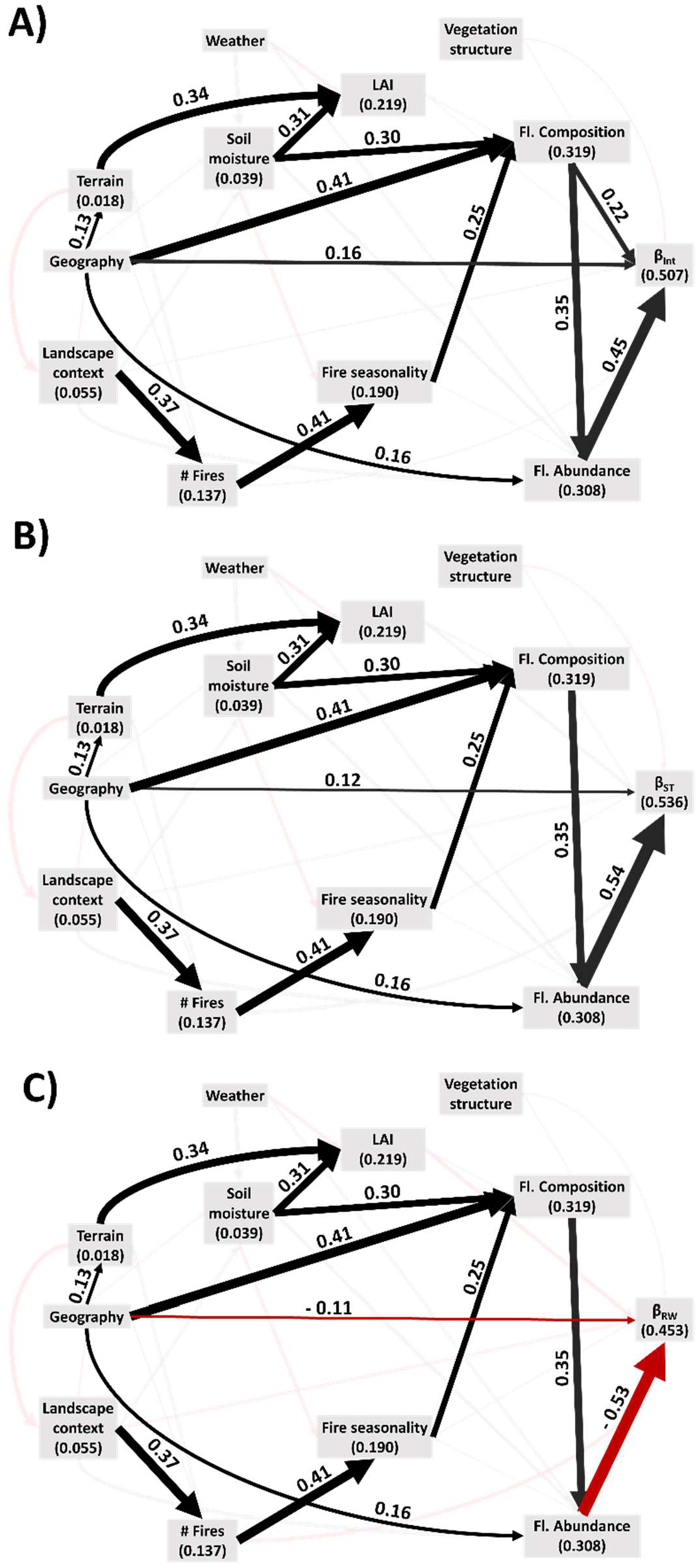
Path analysis for β_Int_, β^ST^, and β_RW_. The names in the boxes refer to site dissimilarities in respect to that particular factor. Positive relationships are portrayed by black lines and indicate parallel trends of both connecting variables (e.g., two sites with similar landscape context have similar number of fires). Negative relationships are shown in red and indicate opposite trends of both connecting variables (e.g., sites with similar soil moisture have different fire seasonality). Bold lines represent significant effects while faded lines represent not significant ones.

Differences in flowering plant composition also had indirect effects on β_Int_ mediated by flower abundance (std. coef. = 0.157). Geographic distance had indirect effects on β_Int_ mediated by flower abundance (std. coef. = 0.070), flowering plant composition (std. coef. = 0.090), and serially though flowering plant composition and flower abundance (std. coef. = 0.065). Additionally, differences in soil moisture, fire seasonality (i.e., growing, and dormant), number of fires since 1990, and spatial context had indirect positive effects on β_Int_ via flowering plant composition and flower abundance (Fig. 3). Differences among site soil moisture and fire seasonality affected β_Int_ via flowering plant composition (soil moisture: std. coef. = 0.065, fire seasonality: std. coef. = 0.056), and serially through flowering plant composition and flower abundance (soil moisture: std. coef. = 0.047, fire seasonality: std. coef. = 0.040). Differences in the number of fires affected β_Int_ via fire seasonality (via flowering plant composition: std. coef. = 0.023, via flowering plant composition and flower abundance: std. coef. = 0.016). Finally, differences in the landscape context altered β_Int_ through the number of fires (direct via flowering plant composition: std. coef. = 0.009, via flower abundance: std. coef. = 0.006). Overall, the SEM for β_Int_ showed that interactions differed most among sites that had dissimilar flower abundance, flowering plant composition, or that are far apart, and that differences in soil moisture, fire seasonality, fire frequency, and landscape context exert an indirect influence on plant-pollinator interactions via flowering plant resources. The model excluding the sites with lower coverage (<0.6) displayed the same patterns, except for the direct path from geographical distance to β_Int_ (P-value = 0.103); the indirect effects of landscape context (mediated only by flower abundance) and soil moisture (negative effect mediated by fire seasonality); and the additional effect of weather (mediated by flower abundance) (Supplementary Material 5 Fig S.5.1, Table S.5.1).

#### Interaction β-diversity due to species turnover (β_ST_)

The best-fit model for β_ST_ (C-statistic = 35.837, df = 68, P-value = 1.000) explained 54% of the variation in interaction β-diversity due to species turnover, supported the hypothesis of flower resources mediated effects (Fig. 2B), and included positive direct effects of differences in flower abundances (std. coef. = 0.538, P-value < 0.001) and geographic distance (std. coef. = 0.117, P-value = 0.039) on β_ST_ (Fig. 3B) (Supplementary Material 3 Table 2). Furthermore, geographic distances had an indirect effect on β_ST_ via flower abundances (std. coef. = 0.083), and serially through flowering plant composition and flower abundance (std. coef. = 0.076). Differences in flowering plant composition had an indirect effect on β_ST_ mediated by flower abundances (std. coef. = 0.187) and differences in soil moisture and fire seasonality had indirect effects on β_ST_ serially mediated by flowering plant composition and flower abundances (soil moisture: std. coef. = 0.055, fire seasonality: std. coef. = 0.047). Divergence in the number of fires since 1990 altered β_ST_ via fire seasonality, flowering plant composition, and flower abundance, serially (std. coef. = 0.019). Finally, differences in the landscape context influenced the number of fires since 1990 and had an indirect effect on β_ST_ following the path of number of fires (std. coef. = 0.007) (Fig. 5). This model indicates that changes in species interactions due to species turnover increased among sites with different flower abundance, or that are located further away. Additionally, sites with greater geographic distance and different landscape contexts, fire frequency, fire seasonality, and/or soil moisture tend to have different flowering plant composition and flower abundance. The model with coverage site > 0.6 included all of the paths but the direct effect of geographic distance (P-value = 0.057) and the effect of soil moisture on flower composition (P-value = 0.132) (Supplementary Material 5). Additionally, the model included a direct effect of flower composition, and indirect effects of weather (mediated via flower abundance) and soil moisture (negative effect via fire seasonality, Fig S.5.1, Table S.5.2).

**Table 2.** Best-fit models for multiple linear regressions of monthly metrics of interaction β-diversity and its partition within preserves against monthly flowering plant characteristics. Levels of significance are noted as:. < 0.1, * <0.05, ** < 0.01, ***<0.001. We present here the results after removing outliers and influential points: 1 for β_Int_, 1 for β_ST_, 4 for β_RW_ (from 68 measurements).

| Metric | Fl. Pl. | Chao 1 | Fl. Pl. | Fl. Pl. | Fl. Pl. | Fl. Pl. | Fl. Pl. | Adj.<br>R<sup>2</sup> |
| --- | --- | --- | --- | --- | --- | --- | --- | --- |
|  | Abundance | Diversity | Richness | Evenness | Abundance | Abundance | Richness |  |
|  |  |  |  |  | x | x | x |  |
|  |  |  |  |  | Chao 1 | Fl. Pl. | Fl. Pl. |  |
|  |  |  |  |  | Diversity | Evenness | Evenness |  |
| $\beta_{Int}$ | -0.040 * | - | 0.019 . | -0.015 | - | -0.031 * | 0.004 | 0.040 |
| $\beta_{ST}$ | -0.077 ** | - | 0.042 | - | - | - | - | 0.093 |
| $\beta_{RW}$ | 0.075 *** | 0.013 | - | - | -0.062 ** | - | - | 0.217 |

#### Interaction β-diversity due to rewiring (β_RW_)

The best-fit SEM for β_RW_ (C-statistic = 37.832, df = 70, P-value = 0.999) described 45% of the variation in β_RW_, supported the hypothesis of flower resources mediated effects (Fig. 2B), and showed that differences in flower abundances (std. coef. = -0.529, P-value < 0.001) and geographic distances reduced β_RW_ (std. coef. = -0.112, P-value = 0.045) (Fig. 3) (Supplementary Material 3 Table 3). Differences in flowering plant composition had an indirect influence on β_RW_ via flower abundance (std. coef. = -0.184). Geographic distances had multiple indirect effects on β_RW_, through flower abundance (std. coef. = -0.082), and flowering plant composition and abundance serially (std. coef. = -0.075). Soil moisture and fire seasonality had negative indirect influences on β_RW_ mediated serially by flowering plant composition and flower abundance (soil moisture: std. coef. = -0.054, fire seasonality: std. coef. = -0.047). Differences in the number of fires affected fire seasonality and, subsequently, β_RW_ (std. coef. = -0.019), following the pathway of fire seasonality. Finally, heterogeneous landscape context altered the number of fires, resulting in a negative effect on β_RW_ via fire and flower resources (std. coef. = -0.007). The β_RW_ SEM shows that species are more likely to interact with different partners in closer sites or sites with similar flower abundances, and that distance, soil moisture, landscape context, and fire frequency, and seasonality drive flower abundance and rewiring via flowering plant composition. The model excluding the sites with poor coverage (coverage < 0.6) was very similar, except for soil moisture, which exerted a positive effect on rewiring mediated serially by fire seasonality, flower composition, and flower abundance (Supplementary Material 5, Fig S.5.1, Table S.5.3).

### Monthly trends of spatial β-diversity

Monthly within-preserve interaction β-diversity and its components were correlated with monthly measures of flowering plant characteristics (Table 2). Β_Int_ had a negative relationship with flower abundance and its interaction with flowering plant evenness. Therefore, preserves with higher flower abundance tend to have less month-to-month variation in plant-pollinator interactions, especially when flower abundances are even among plant species (Table 2). The best-fit model explained 3.7% of the variance in β_Int_. β_ST_ decreased with flower abundance which indicated that plant-pollinator interactions are more uniform across sites within preserves that have more flowers (Table 2). The best-fit model explained 13.7% of the variance in β_ST_. β_RW_ had a positive relationship with flower abundance but a negative relationship with the interaction between flower abundance and flowering plant diversity measured by the Chao 1 index (Table 2). Hence, plant and pollinator species tended to conserve their interactions across months in preserves with lower flower abundances, especially when they also have high flowering plant diversity. The best-fit model explained 21.7% the variance of β_RW_. The models including only the preserves with coverage > 0.6 per month included all the aforementioned trends as well as some additional effects of flower diversity (Chao 1 index, and richness) (Supplementary Material 5, Tables S.5.4, and S.5.5).

## DISCUSSION

Plant-pollinator interactions in Florida savannas are highly variable across space and time, similar to other plant-pollinator systems (CaraDonna et al. 2017; Simanonok and Burkle 2014; Trøjelsgaard et al. 2015; Souza et al. 2021). Spatio-temporal interaction β-diversity is largely driven by species turnover. The high contribution of species turnover to spatial interaction β-diversity is comparable to trends observed in other systems (Trøjelsgaard et al. 2015; Simanonok and Burkle 2014) and suggests that rewiring of plant-pollinator interactions is not very common across space, possibly due to spatial abiotic gradients (Simanonok and Burkle 2014; Trøjelsgaard et al. 2015) or dispersal boundaries (Trøjelsgaard et al. 2015).

### Path analysis: biogeographical and ecological drivers

Flower resources exerted the largest effect on plant-pollinator interaction β-diversity (and its components), and mediated the effects of fire regime, landscape connectivity and soil moisture providing support for the flower resources mediation hypothesis (Fig. 2B). Flower abundance was the most important direct driver of interaction β-diversity and its components (i.e., species turnover and interaction rewiring), with path standardized coefficients two to five times larger than those of other significant drivers. Flower abundance has been related to temporal interaction β-diversity and rewiring in another system (CaraDonna et al. 2017) and numerous studies have suggested that flower abundance may modulate the richness of pollinators (Stang, Klinkhamer, and Meijden 2006; Kuussaari et al. 2007; Hall and Ascher 2011; Atwater 2013; Ponisio et al. 2016; Lucas et al. 2017), and plant-pollinator interactions (Ponisio et al. 2016). Our results further suggest that higher flower abundance may not only increase the richness of pollinator species, plant-pollinator assemblages, and interactions, but also that sites in the same system with many blooming flowering plants have similar plant-pollinator composition and interactions. It is possible that sites with many flowering plants not only include more species but also similar ones (e.g., specialists), which are absent in sites with low numbers of flowering plants.

Flower abundance had a negative effect on rewiring, which suggests that sites with similar abundances of blooming flowering plants exhibit frequent interaction rewiring. Rewiring is the mechanism by which species switch partners across sites despite homogeneous partner availability (Poisot et al. 2012). In our system, flowering plant composition altered rewiring via flower abundance, but did not exert a direct effect indicating the presence of a flowering plant does not guarantee its participation in pollination networks. This result is consistent with previous studies that reported joint effects of species presences and abundances on rewiring (CaraDonna et al. 2017). Rewiring can further depend on additional factors including species relative abundances (Trøjelsgaard et al. 2015; CaraDonna et al. 2017), specialization (Trøjelsgaard et al. 2015), and competition (Simanonok and Burkle 2014; Trøjelsgaard et al. 2015; CaraDonna et al. 2017). Pine savannas with many blooming flowering plants may display frequent interaction rewiring depending on species evenness, specialization, or pollinator competition, whereas savannas with few blooming flowering plants are likely to have lower interaction rewiring.

Geographical distances had consistent albeit smaller effects on interaction β-diversity than flower abundances (i.e., path standardized coefficients three to five times smaller). Geographically closer sites had more similar plant-pollinator interactions and assemblages, but higher levels of rewiring. The increase in interaction β-diversity and the contribution of species turnover with spatial distance reflects the distance decay of species and interactions, and may be the result of differences among habitats or environmental factors, as well as connectivity limitations (Simanonok and Burkle 2014; Trøjelsgaard et al. 2015; de Manincor et al. 2020; Souza et al. 2021). Lower species turnover among closer sites allows, but do not ensure, interaction rewiring. Rewiring among closer sites is more common, but depends on species relative abundances, specialization, and/or competition.

Flowering plant composition altered interaction β-diversity and its components via flower abundance, exerting a similar effect to that of geographic distances (i.e., similar path standardized coefficients). Additionally, flowering plant composition exerted a larger direct effect on β_Int_ than that of geographical distances (i.e., path standardized coefficient about 1.4 times higher). Sites with similar species of flowering plants tended to have similar plant-pollinator assemblages and interactions, but lower rewiring. This pattern suggests that sandhill pollinators are at least moderately specialized, as suggested for other systems (Blüthgen et al. 2007; Dormann et al. 2009). Sites with similar flowering plants attract similar pollinators, include similar plant-pollinator interactions, and enable interaction rewiring. Flowering plant composition was also a key mediator, modulating the effects of soil moisture, fire, and landscape structure.

Soil moisture, fire frequency and seasonality, and landscape context had indirect effects on interaction β-diversity and its partitions of smaller magnitude (|path standardized coefficients| < 0.01). Sites with similar soil moisture had similar flowering plant composition and flower abundances. Consequently, soil moisture had an indirect effect on plant-pollinator interaction β-diversity and its components, by which sites with similar soil moisture included similar species assemblages and interactions but frequent rewiring. The effect of soil moisture driving plant composition is well known in pine savannas (Platt 2007). Fire frequency and seasonality were also important variables driving plant composition and, ultimately, plant-pollinator interactions. Sites with similar fire frequencies were more likely to exhibit similar fire seasonality, and plant composition, which subsequently led to similar plant-pollinator assemblages and interactions but enabled frequent rewiring. Fire has been shown to alter multiple plant characteristics in pine savannas, including phenology, abundance, number of flowers per plant, and peak flowering times (Platt, Evans, and Davis 1988). Fire was also driven by the spatial context of the sites. Sites with similar surrounding habitats had similar fire frequencies, and seasonality, driving plant-pollinator interactions via plant composition. Landscape fragmentation has been demonstrated to limit prescribed fires in pine savannas (Duncan and Schmalzer 2004; Duncan, Shao, and Adrian 2009).

### Monthly trends of spatial β-diversity

Spatial interaction β-diversity among sites within the same preserve changed across the flowering season. Interaction β-diversity and its partition due to species turnover decreased with monthly flower abundance indicating that plant-pollinator composition and interactions are more uniform across space when blooming flowering plants are more abundant. In contrast, interaction β-diversity due to rewiring increased when blooming flowering plants were more abundant. Flower abundance increases the available resources for pollinators (Heithaus 1974; Lucas et al. 2017), expands niche space (Heithaus 1974; Lara-Romero et al. 2016), and decreases competition among pollinator species (Escobedo-Kenefic et al. 2020), fostering rich and abundant pollinator assemblages (Stang, Klinkhamer, and Meijden 2006; Kuussaari et al. 2007; Hall and Ascher 2011; Ponisio et al. 2016; Lucas et al. 2017) and plant-pollinator networks (Ponisio et al. 2016). High flower abundances within preserves may decrease the spatial segregation of species, and promote pollinator connectivity, due to the reduction in competition, as well as the increases of pollinator abundance and richness. In the pine savanna ecosystem, high flower abundances are especially common in fall, when composite species bloom (Asteraceae, Magnoliopsida: Euasterids) (Platt, Evans, and Davis 1988; Heuberger and Putz 2003; Hall and Ascher 2011) suggesting that pollinator networks may be more homogeneous across sites during this time. High plant-pollinator homogeneity reduced interaction β-diversity as well as the contribution of species turnover but enabled more opportunities for rewiring. Therefore, sites with similar flower abundances fostered similar annual plant-pollinator interactions and compositions while enabling rewiring, and preserves with high flowering abundances foster more uniform monthly interactions and compositions but higher rewiring across sites.

Interaction β-diversity and rewiring varied with the statistical interaction between flower abundance and other measures of floral resources. Interaction β-diversity decreased with the statistical interaction between flower abundance and plant evenness indicating that plant-pollinator interactions were similar within preserves when blooming flowering plants were abundant and even. Uniform plant abundances within preserves promote uniform species assemblages and decrease pollinator competition (Escobedo-Kenefic et al. 2020), promoting homogenous interactions. Additionally, plant evenness homogenizes relative abundances, which reduces rewiring (CaraDonna et al. 2017; Trøjelsgaard et al. 2015). Thus, plant communities that are species-rich or have high evenness may simultaneously limit the contributions of species turnover and interaction rewiring, promoting low interaction β-diversity across space.

Finally, rewiring decreased with the statistical interaction between flower abundance and diversity, showing that rewiring among sites within a preserve was more common when preserves included more abundant but less species-rich flowering plant assemblages. Rewiring was, however, not significantly affected by flowering plant evenness, suggesting that the interactive effect of flower abundance and flowering plant diversity is independent from flowering plant distributions. This effect may be explained by pollinator preferences and partner availability such that in diverse plant communities, pollinators are more likely to find their preferred resources and, consequently, less likely to switch floral partners. Conversely, in species-poor communities, pollinators are less likely to encounter their preferred resources and interact with different plants depending on the local context.

### Management implications

Fragmentation and fire suppression are seen as the biggest threats to pine sandhills (e.g., Heuberger and Putz 2003; Van Lear et al. 2005; Platt 2007; Atwater 2013) and we show that they can synergistically influence plant-pollinator networks. We found that fragmented landscapes are also more likely to be fire suppressed, a finding that is consistent with previous work (Duncan and Schmalzer 2004; Duncan, Shao, and Adrian 2009), ultimately altering plant composition and plant-pollinator interactions. The threats of fire suppression and habitat fragmentation are also likely to increase in the near future, driven by current development plans (Boda 2018). It is essential to protect extant sandhills, promote corridors, and practice good fire management to preserve pollination systems, which are pivotal to natural ecosystems and human agriculture (Garibaldi et al. 2013).

## Conclusion

Sandhill plant-pollinator interactions are highly variable across space and time. These differences are mainly due to changes in plant-pollinator interactions due to species turnover, especially at smaller spatial and temporal scales (i.e., surveys, sites, and months). Our findings suggest that sandhill plant-pollinator interactions are spatially and temporally structured, possibly due to the combination of environmental gradients and habitat fragmentation. When we specifically tested the ecological and biogeographical causality pathways, we found that interaction β-diversity in sandhill plant-pollinator networks is mainly explained by local flowering plant resources, as well as geographic distance. However, interaction β-diversity and its components are also indirectly altered by fire regimes, soil moisture, and landscape context. Soil moisture and the ecosystem connectivity-fire relationship underlie flowering plant diversity and abundance in pine sandhills. Sandhills with higher soil moisture tend to have lower flower abundance. Additionally, sandhills that are surrounded by other natural sandhills tend to have a fire regime that better promotes high flower abundances, likely due to the increased fire constraints managers face in more fragmented landscapes. While soil moisture is not within our control, managers can promote landscape connectivity and maintain appropriate fire regimes on sandhills, fostering richer, more complex, and robust plant-pollinator communities. The conservation of the remaining pine sandhills is essential, since they are critically imperiled habitats that occupy 2-3% of their historical range and are currently threatened by regional development plans. By conserving sandhill plant-pollinator interactions, we can maintain sandhill structure, functioning and ecosystem services, including the pollination of agricultural crops.

