## Supplementary materials for "Plant-mediated effects of fire and fragmentation drive plant-pollinator interaction β-diversity in fire-dependent pine savannas"

**Supplementary material. Index**

### Supplementary material 1. Field sites

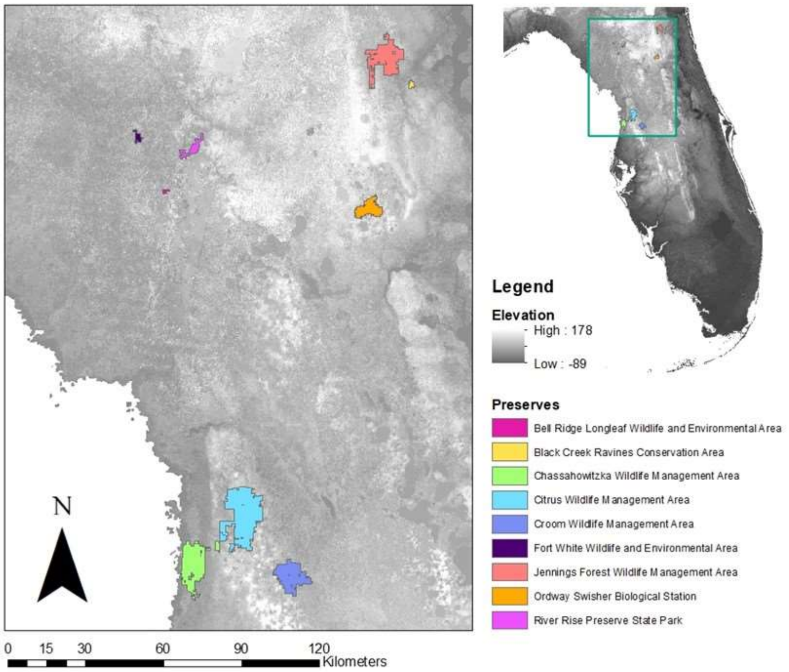

Figure S.1. Study preserves for plant-visitor interactions. All preserves are located within Timucua and Seminole native territories. Additionally, Chassahowitzka Wildlife Management Area and the Croom tract within Withlacoochee State Forest are partially located within Tocobaga native territory (Temprano 2015).

### Supplementary material 2. Insect guides

Table S.2. List of references for insect identification

| Taxa | Reference |
|---|---|
| **INSECTA** | Bartlett, Troy. "BugGuide. Net: Identification, Images, & Information For Insects, Spiders & Their Kin For the United States & Canada." (2003). |
| | Pickering, John. "Discover life." University of Georgia Athens (URL: http://discoverlife. org/) (2009). |
| **Coleoptera** | Epler, J. H. "The Water Beetles of Florida-an identification manual for the families Chrysomelidae." Curculionidae, Dryopidae, Elmidae, Gyrinidae, Haliplidae, Helophoridae, Hydraenidae, Hydrochidae, Hydrophilidae, Noteridae, Psephenidae, Ptilodactylidae and Scirtidae. Florida Department of Environmental Protection, Tallahassee, FL 399 (2010). |
| *Belotus sp.* | Ramsdale, Alistair S. "Family 64. Cantharidae", in *American beetles, volume II: Polyphaga: Scarabaeoidea through Curculionoidea,* edited by Ross H. Arnett Jr., Michael C. Thomas, Paul E. Skelley, and J. Howard Frank. CRC Press, 2002. |
| Carabidae | Bousquet, Yves. Illustrated Identification Guide to Adults and Larvae of Northeastern North American Ground Beetles (Coleoptera, Carabidae) (Pensoft Series Faunistica, No. 90). Pensoft Publishers, 2010. |
| *Collops sp.* | Fall, H. C. "A review of the North American species of *Collops* (Col.)." Journal of the New York Entomological Society 20, no. 4 (1912): 249-274. |
| | Mayor, Adriean J. "Family 74. Melyridae", in *American beetles, volume II: Polyphaga: Scarabaeoidea through Curculionoidea,* edited by Ross H. Arnett Jr., Michael C. Thomas, Paul E. Skelley, and J. Howard Frank. CRC Press, 2002. |
| *Epicauta sp.* | Pinto, John D. and Marco A. Bologna "Family 111. Meloidae", in *American beetles, volume II: Polyphaga: Scarabaeoidea through Curculionoidea,* edited by Ross H. Arnett Jr., Michael C. Thomas, Paul E. Skelley, and J. Howard Frank. CRC Press, 2002. |
| Mordellidae | Liljeblad, Emil. "Monograph of the family Mordellidae (Coleoptera) of North America, north of Mexico." (1945). |
| *Nemognatha sp.* | Enns, Wilbur R. "A revision of the genera *Nemognatha*, *Zonitis*, and *Pseudozonitis* (Coleoptera, Meloidae) in America north of Mexico, with a proposed new genus." PhD diss., University of Kansas, Entomology, 1955. |
| **Diptera** | McAlpine, J. F., B. V. Peterson, G. E. Shewell, H. J. Teskey, J. R. Vockeroth, and D. M. Wood. Manual of Nearctic Diptera. Volume 1. No. 27. 1981. |
| | McAlpine, J. F., B. V. Peterson, G. E. Shewell, H. J. Teskey, J. R. Vockeroth, and D. M. Wood. Manual of Nearctic Diptera. Volume 2. No. 28. 1992. |

**Hymenoptera**

*Ammophila sp.* Menke, A. S. "A new subgenus of *Ammophila* from the Neotropical Region (Hymenoptera: Sphecidae)." The Canadian Entomologist 96, no. 6 (1964): 874-883.

Antophila Pascarella, J. B. and H. Glenn. "The Bees of Florida". (2021).

*Bembecinus sp.* Bohart, Richard Mitchell. "A review of *Bembecinus* (Hymenoptera: Sphecidae: Stizini) in North and Central America." Entomological Society of Washington (USA) (1996).

*Bembix sp.* Bohart, Richard Mitchell, and Donald S. Horning. "California bembicine sand wasps." (1971): 1.

Evans, Howard E., and Robert W. Matthews. "North American *Bembix*, a revised key and suggested grouping." Annals of the Entomological Society of America 61, no. 5 (1968): 1284-1299.

*Bycirtes sp.* Bohart, Richard M. "A review of the genus *Bicyrtes* (Hymenoptera: Sphecidae, Nyssoninae, Bembicini)." Insecta Mundi (1996): 4.

*Cerceris sp.* Scullen, Herman A. "Review of the genus *Cerceris* in America north of Mexico (Hymenoptera: Sphecidae)." Proceedings of the United States National Museum (1965).

*Crabro sp.* Bohart, Richard M. "A review of the Nearctic species of *Crabro* (Hymenoptera: Sphecidae)." Transactions of the American Entomological Society (1890-) 102, no. 2 (1976): 229-287.

*Dasymutilla sp.* Manley, Donald G., Kevin A. Williams, and James P. Pitts. "Keys to nearctic velvet ants of the genus *Dasymutilla* Ashmead (Hymenoptera: Mutillidae), with notes on taxonomic changes since Krombein (1979)." Proceedings of the Entomological Society of Washington 122, no. 2 (2020): 335-414.

*Ephuta sp.* Schuster, R. M. "A revision of the genus *Ephuta* (Mutillidae) in America north of Mexico." Journal of the New York Entomological Society 59, no. 1 (1951): 1-43.

Schuster, R. M. "A Revision of the Genus *Ephuta* (Mutillidae) in America north of Mexico." Journal of the New York Entomological Society 64 (1956): 7-84.

*Lasioglossum sp.* Gibbs, Jason. "Revision of the metallic species of *Lasioglossum* (*Dialictus*) in Canada (Hymenoptera, Halictidae, Halictini)." Zootaxa 2591, no. 1 (2010): 1-382.

*Liris sp.* Krombein, Karl V., and Sandra Shanks Gingras. "Revision of North American *Liris* Fabricius (Hymenoptera: Sphecoidea: Larridae)." (1984).

Mutilidae Manley, D. G., and J. P. Pitts. "A key to genera and subgenera of Mutillidae (Hymenoptera) in America north of Mexico with description of a new genus." Journal of Hymenoptera Research 11, no. 1 (2002): 72-100.

*Myzinum sp.* Kimsey, Lynn S. "Taxonomic purgatory: Sorting out the wasp genus *Myzinum* Latreille in North America (Hymenoptera, Tiphiidae, Myzininae)." Zootaxa 2224, no. 1 (2009): 30-50.

*Paratiphia sp.* Allen, Harry W. "A monographic study of the genus *Paratiphia*." Transactions of the American Entomological Society (1890-) 94, no. 1 (1968): 25-109.

Philanthinae Bohart, Richard Mitchell, and E. E. Grissell. "California wasps of the subfamily Philanthinae (Hymenoptera: Sphecidae)." (1975).

| | |
|---|---|
| Pompilidae | Kurczewski, Frank E., and G. B. Edwards. "Hosts, nesting behavior, and ecology of some North American spider wasps (Hymenoptera: Pompilidae)." Southeastern Naturalist 11, no. m4 (2012): 1-71. |
| | Evans, Howard E. "A taxonomic study of the Nearctic spider wasps belonging to the tribe Pompilini (Hymenoptera: Pompilidae). Part I." Transactions of the American Entomological Society (1890-) 75, no. 3/4 (1949): 133-270. |
| | Evans, Howard E. "A taxonomic study of the Nearctic spider wasps belonging to the tribe Pompilini (Hymenoptera: Pompilidae). Part II: Genus *Anoplius* Dufour." Transactions of the American Entomological Society (1890-) 76, no. 4 (1950): 207-361. |
| Sphecidae | Bohart, Richard M., Richard Mitchell Bohart, and Arnold S. Menke. Sphecid wasps of the world: a generic revision. Univ of California Press, 1976. |
| *Tachytes sp.* | Bohart, Richard M. "A key to the genus *Tachytes* in America north of Mexico with descriptions of three new species." Proc. Entomol. Soc. Wash 96, no. 2 (1994): 342-349. |
| Tiphiinae | Allen, Harry W. "A revision of the Tiphiinae (Hymenoptera: Tiphiidae) of eastern North America." Transactions of the American Entomological Society (1890-) 92, no. 2 (1966): 231-356. |
| Vespidae | Buck, Matthias, Stephen A. Marshall, and David KB Cheung. "Identification Atlas of the Vespidae (Hymenoptera, Aculeata) of the northeastern Nearctic region." Canadian journal of arthropod identification 5, no. 1 (2008): 1-492. |
| *Zethus sp.* | Porter, Charles C. "Ecology and Taxonomy of Lower Rio Grande Valley *Zethus*." Florida Entomologist (1978): 159-167. |
| **Lepidoptera** | Glassberg, Jeffrey, Marc C. Minno, and John V. Calhoun. Butterflies through binoculars. Oxford University Press, 2000. |
| | Heppner, J. B. "Lepidoptera of Florida. Part 1. Introduction and catalog." Arthropods of Florida and neighboring land areas 17 (2003): 670. |

### Supplementary material 3. Coefficients of permutation-based path analysis models

#### Plant-pollinator interactions ($\beta_{Int}$)

Table S.3.1. Coefficients path analysis for plant-pollinator interactions ($\beta_{Int}$). We used permutation-based path analysis (Fourtune et al. 2018). Levels of significance are noted as: . < 0.1, * <0.05, ** < 0.01, ***<0.001.

| Path | Std. Estimate | Lower CI | Upper CI |
|---|---|---|---|
| $\beta_{Int}$ ~ flowering plant abundance dissimilarity | 0.454 *** | 0.357 | 0.551 |
| $\beta_{Int}$ ~ flowering plant composition dissimilarity | 0.220 ** | 0.118 | 0.322 |
| $\beta_{Int}$ ~ vegetation structure dissimilarity | -0.064 | -0.147 | 0.020 |
| $\beta_{Int}$ ~ LAI diss. | 0.076 | -0.009 | 0.161 |
| $\beta_{Int}$ ~ number fires dissimilarity | 0.090 | 0.001 | 0.179 |
| $\beta_{Int}$ ~ landscape context dissimilarity | 0.085 | -0.005 | 0.175 |
| $\beta_{Int}$ ~ geographic distance | 0.162 ** | 0.070 | 0.253 |
| Fl. plant abund. diss. ~ Fl. plant comp. diss. | 0.347 *** | 0.234 | 0.460 |
| Fl. plant abund. diss. ~ fire seasonality (2-S) diss. | 0.065 | -0.040 | 0.169 |
| Fl. plant abund. diss. ~ soil moisture diss. | 0.163 . | 0.056 | 0.271 |
| Fl. plant abund. diss. ~ land context diss. | 0.159 | 0.060 | 0.257 |
| Fl. plant abund. diss. ~ weather diss. | 0.141 . | 0.042 | 0.239 |
| Fl. plant abund. diss. ~ geographic dist. | 0.155 * | 0.046 | 0.264 |
| Fl. plant comp. diss. ~ veg. structure diss. | 0.087 | -0.010 | 0.184 |
| Fl. plant comp. diss. ~ LAI diss. | 0.105 | 0.004 | 0.207 |
| Fl. plant comp. diss. ~ fire seasonality (2-S) diss. | 0.253 * | 0.157 | 0.350 |
| Fl. plant comp. diss. ~ soil moisture diss. | 0.296 ** | 0.196 | 0.397 |
| Fl. plant comp. diss. ~ geographic dist. | 0.410 *** | 0.323 | 0.496 |
| LAI diss. ~ soil moisture diss. | 0.316 * | 0.214 | 0.417 |
| LAI diss. ~ terrain diss. | 0.344 * | 0.245 | 0.442 |
| LAI diss. ~ weather diss. | -0.140 | -0.244 | -0.035 |
| fire seas. (2-S) diss. ~ number fires diss. | 0.410 ** | 0.312 | 0.507 |
| fire seas. (2-S) diss. ~ soil moisture diss. | -0.149 | -0.254 | -0.044 |
| number fires diss. ~ land context diss. | 0.373 ** | 0.268 | 0.478 |
| number fires diss. ~ terrain diss. | 0.143 | 0.031 | 0.255 |
| number fires diss. ~ geographic dist. | 0.064 | -0.047 | 0.174 |
| soil moisture diss. ~ weather diss. | 0.182 | 0.069 | 0.295 |
| soil moisture diss. ~ geographic dist. | -0.090 . | -0.205 | 0.025 |
| land context diss. ~ terrain diss. | -0.216 . | -0.329 | -0.103 |
| land context diss. ~ geographic dist. | -0.068 | -0.183 | 0.048 |
| terrain diss. ~ geographic dist. | 0.133 * | 0.018 | 0.248 |

**$\beta$-diversity due to species turnover ($\beta_{ST}$)**

Table S.3.2. Coefficients path analysis for interaction $\beta$-diversity due to species turnover ($\beta_{ST}$). We used permutation-based path analysis (Fourtune et al. 2018). Levels of significance are noted as: . < 0.1, * <0.05, ** < 0.01, ***<0.001.

| Path | Std. Estimate | Lower CI | Upper CI |
|---|---|---|---|
| $\beta_{ST}$ ~ flowering plant abundance dissimilarity | 0.538 *** | 0.453 | 0.622 |
| $\beta_{ST}$ ~ flowering plant composition dissimilarity | 0.100 | 0.000 | 0.199 |
| $\beta_{ST}$ ~ vegetation structure dissimilarity | -0.096 | -0.176 | -0.015 |
| $\beta_{ST}$ ~ LAI dissimilarity | 0.165 . | 0.083 | 0.248 |
| $\beta_{ST}$ ~ fire seasonality (2-S) diss. | -0.069 | -0.159 | 0.021 |
| $\beta_{ST}$ ~ number fires dissimilarity | 0.160 | 0.067 | 0.253 |
| $\beta_{ST}$ ~ landscape context dissimilarity | 0.116 | 0.029 | 0.204 |
| $\beta_{ST}$ ~ geographic distance | 0.117 * | 0.028 | 0.206 |
| Fl. plant abund. diss. ~ Fl. plant comp. diss. | 0.347 *** | 0.234 | 0.460 |
| Fl. plant abund. diss. ~ fire seasonality (2-S) diss. | 0.065 | -0.040 | 0.169 |
| Fl. plant abund. diss. ~ soil moisture diss. | 0.163 . | 0.056 | 0.271 |
| Fl. plant abund. diss. ~ land context diss. | 0.159 . | 0.060 | 0.257 |
| Fl. plant abund. diss. ~ weather diss. | 0.141 . | 0.042 | 0.239 |
| Fl. plant abund. diss. ~ geographic dist. | 0.155 * | 0.046 | 0.264 |
| Fl. plant comp. diss. ~ veg. structure diss. | 0.087 | -0.010 | 0.184 |
| Fl. plant comp. diss. ~ LAI diss. | 0.105 | 0.004 | 0.207 |
| Fl. plant comp. diss. ~ fire seasonality (2-S) diss. | 0.253 * | 0.157 | 0.350 |
| Fl. plant comp. diss. ~ soil moisture diss. | 0.296 ** | 0.196 | 0.397 |
| Fl. plant comp. diss. ~ geographic dist. | 0.410 *** | 0.323 | 0.496 |
| LAI diss. ~ soil moisture diss. | 0.316 * | 0.214 | 0.417 |
| LAI diss. ~ terrain diss. | 0.344 * | 0.245 | 0.442 |
| LAI diss. ~ weather diss. | -0.140 | -0.244 | -0.035 |
| fire seas. (2-S) diss. ~ number fires diss. | 0.410 ** | 0.312 | 0.507 |
| fire seas. (2-S) diss. ~ soil moisture diss. | -0.149 | -0.254 | -0.044 |
| number fires diss. ~ land context diss. | 0.373 * | 0.268 | 0.478 |
| number fires diss. ~ terrain diss. | 0.143 | 0.031 | 0.255 |
| number fires diss. ~ geographic dist. | 0.064 | -0.047 | 0.174 |
| soil moisture diss. ~ weather diss. | 0.182 | 0.069 | 0.295 |
| soil moisture diss. ~ geographic dist. | -0.090 . | -0.205 | 0.025 |
| land context diss. ~ terrain diss. | -0.216 . | -0.329 | -0.103 |
| land context diss. ~ geographic dist. | -0.068 | -0.183 | 0.048 |
| terrain diss. ~ geographic dist. | 0.133 * | 0.018 | 0.248 |

### Rewiring ($\beta_{RW}$)

Table S.3.3. Coefficients path analysis for rewiring ($\beta_{RW}$). We used permutation-based path analysis (Fourtune et al. 2018). Levels of significance are noted as: . < 0.1, * <0.05, ** < 0.01, ***<0.001.

| Path | Std. Estimate | Lower CI | Upper CI |
|---|---|---|---|
| $\beta_{RW}$ ~ flowering plant abundance dissimilarity | -0.529 *** | -0.612 | -0.446 |
| $\beta_{RW}$ ~ vegetation structure dissimilarity | 0.090 | 0.003 | 0.177 |
| $\beta_{RW}$ ~ LAI dissimilarity | -0.181 . | -0.268 | -0.093 |
| $\beta_{RW}$ ~ fire seasonality (2-S) dissimilarity | 0.073 | -0.023 | 0.169 |
| $\beta_{RW}$ ~ number fires dissimilarity | -0.163 . | -0.264 | -0.062 |
| $\beta_{RW}$ ~ landscape context distance | -0.110 | -0.205 | -0.015 |
| $\beta_{RW}$ ~ geographic distance | -0.112 * | -0.204 | -0.021 |
| Fl. plant abund. diss. ~ Fl. plant comp. diss. | 0.347 *** | 0.234 | 0.460 |
| Fl. plant abund. diss. ~ fire seasonality (2-S) diss. | 0.065 | -0.040 | 0.169 |
| Fl. plant abund. diss. ~ soil moisture diss. | 0.163 . | 0.056 | 0.271 |
| Fl. plant abund. diss. ~ land context diss. | 0.159 . | 0.060 | 0.257 |
| Fl. plant abund. diss. ~ weather diss. | 0.141 . | 0.042 | 0.239 |
| Fl. plant abund. diss. ~ geographic dist. | 0.155 * | 0.046 | 0.264 |
| Fl. plant comp. diss. ~ veg. structure diss. | 0.087 | -0.010 | 0.184 |
| Fl. plant comp. diss. ~ LAI diss. | 0.105 | 0.004 | 0.207 |
| Fl. plant comp. diss. ~ fire seasonality (2-S) diss. | 0.253 * | 0.157 | 0.350 |
| Fl. plant comp. diss. ~ soil moisture diss. | 0.296 * | 0.196 | 0.397 |
| Fl. plant comp. diss. ~ geographic dist. | 0.410 *** | 0.323 | 0.496 |
| LAI diss. ~ soil moisture diss. | 0.316 * | 0.214 | 0.417 |
| LAI diss. ~ terrain diss. | 0.344 ** | 0.245 | 0.442 |
| LAI diss. ~ weather diss. | -0.140 | -0.244 | -0.035 |
| fire seas. (2-S) diss. ~ number fires diss. | 0.410 ** | 0.312 | 0.507 |
| fire seas. (2-S) diss. ~ soil moisture diss. | -0.149 | -0.254 | -0.044 |
| number fires diss. ~ land context diss. | 0.373 * | 0.268 | 0.478 |
| number fires diss. ~ terrain diss. | 0.143 | 0.031 | 0.255 |
| number fires diss. ~ geographic dist. | 0.064 | -0.047 | 0.174 |
| soil moisture diss. ~ weather diss. | 0.182 | 0.069 | 0.295 |
| soil moisture diss. ~ geographic dist. | -0.090 . | -0.205 | 0.025 |
| land context diss. ~ terrain diss. | -0.216 . | -0.329 | -0.103 |
| land context diss. ~ geographic dist. | -0.068 | -0.183 | 0.048 |
| terrain diss. ~ geographic dist. | 0.133 * | 0.018 | 0.248 |

### Supplementary material 4. Monthly models of spatial β-diversity with outliers

Table S.4.1. Best-fit models for multiple linear regressions of monthly metrics of interaction β-diversity and its partition within preserves against monthly flowering plant characteristics. Raw models with outliers. Levels of significance are noted as: . < 0.1, * <0.05, ** < 0.01, ***<0.001.

| | Fl. Pl. Abundance | Chao 1 Diversity | Fl. Pl. Richness | Fl. Pl. Evenness | Fl. Pl. Abundance x Chao 1 Diversity | Fl. Pl. Abundance x Fl. Pl. Evenness | Fl. Pl. Richness x Fl. Pl. Evenness | Adj. $R^2$ |
|---|---|---|---|---|---|---|---|---|
| $\beta_{Int}$ | -0.041 * | - | 0.031 ** | -0.018 | - | -0.030 . | 0.016 . | 0.097 |
| $\beta_{ST}$ | -0.074 ** | - | 0.038 | - | - | - | - | 0.086 |
| $\beta_{RW}$ | 0.067 *** | 0.001 | - | - | -0.048 * | - | - | 0.157 |

**Supplementary material 5. Permutation-based path analysis models including samples with coverage > 0.6**

Interaction sampling coverage, the probability of occurrence of interactions within the network samples, can affect the estimation of interaction networks and, subsequently, derived metrics such as β-diversity. Our samples for the path analyses (sites for all months) had a median coverage of 0.66 and standard deviation of 0.11; those for the monthly trends of β-diversity (preserves per month) had a median coverage of 0.63 with standard deviation of 0.20. We re-analyzed the models using only those samples (sites over all months for the path analyses, and preserves per each month for the monthly trends of spatial β-diversity) that had a coverage > 0.6. While interaction sample coverage has not been extensively studied, some classic plant-pollinator networks have similar coverages (e.g., Zackenberg, Olesen et al. 2008; Jordano 2016).

### PATH ANALYSIS: BIOGEOGRAPHICAL AND ECOLOGICAL DRIVERS

Our path analyses included 19 of the original 24 sites (i.e., all but WIN1, WIS1, BR2, RR1, and JE2).

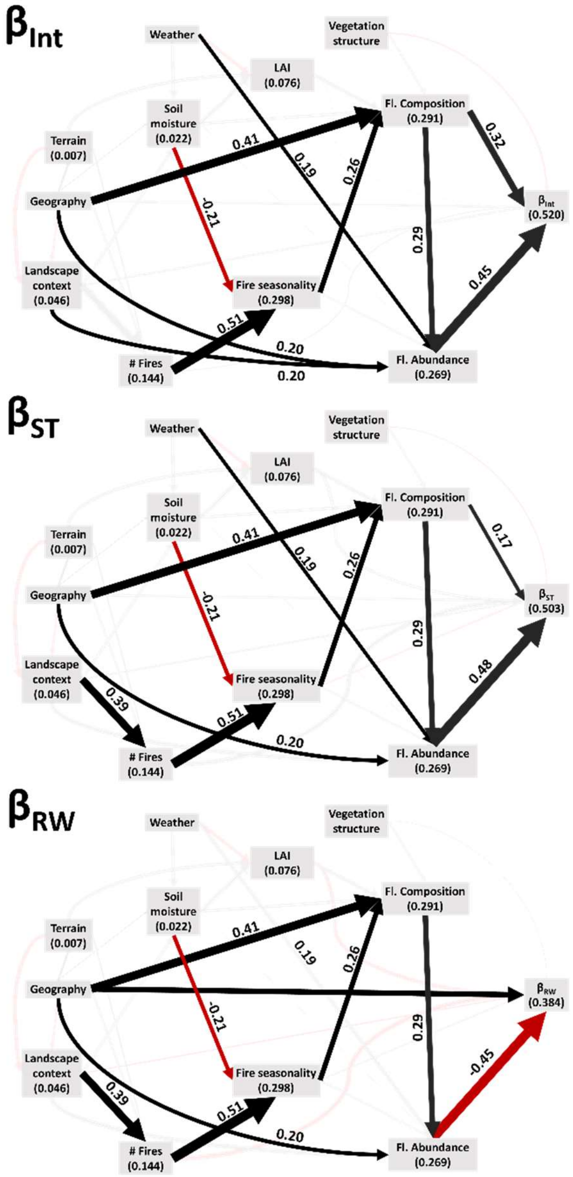

Figure S.5. 1. Path analysis for $\beta_{Int}$, $\beta_{ST}$, and $\beta_{RW}$ including sites with coverage > 0.6. The names in the boxes refer to site dissimilarities in respect to that particular factor. Positive relationships are portrayed by black lines and indicate parallel trends of both connecting variables (e.g., two sites with similar landscape context have similar number of fires). Negative relationships are shown in red and indicate opposite trends of both connecting variables (e.g., sites with similar soil moisture have different fire seasonality). Bold lines represent significant effects while faded lines represent not significant ones.

### Plant-pollinator interactions ($\beta_{Int}$)

Table S.5.1. Coefficients path analysis for plant-pollinator interactions ($\beta_{Int}$). We used permutation-based path analysis (Fourtune et al. 2018). Levels of significance are noted as: . < 0.1, * <0.05, ** < 0.01, ***<0.001.

| Path | Std. Estimate | Lower CI | Upper CI |
|---|---|---|---|
| $\beta_{Int}$ ~ flowering plant abundance dissimilarity | 0.452 *** | 0.341 | 0.563 |
| $\beta_{Int}$ ~ flowering plant composition dissimilarity | 0.316 *** | 0.194 | 0.438 |
| $\beta_{Int}$ ~ vegetation structure dissimilarity | -0.082 | -0.186 | 0.023 |
| $\beta_{Int}$ ~ LAI diss. | 0.004 | -0.102 | 0.110 |
| $\beta_{Int}$ ~ number fires dissimilarity | 0.041 | -0.073 | 0.154 |
| $\beta_{Int}$ ~ land context diss. | 0.099 | -0.016 | 0.214 |
| $\beta_{Int}$ ~ geographic distance | 0.096 | -0.022 | 0.214 |
| Fl. plant abund. diss. ~ Fl. plant comp. diss. | 0.286 *** | 0.142 | 0.430 |
| Fl. plant abund. diss. ~ fire seasonality (2-S) diss. | 0.105 | -0.033 | 0.244 |
| Fl. plant abund. diss. ~ soil moisture diss. | 0.009 | -0.126 | 0.145 |
| Fl. plant abund. diss. ~ land context diss. | 0.201 * | 0.071 | 0.331 |
| Fl. plant abund. diss. ~ weather diss. | 0.188 * | 0.061 | 0.315 |
| Fl. plant abund. diss. ~ geographic dist. | 0.201 * | 0.061 | 0.340 |
| Fl. plant comp. diss. ~ veg. structure diss. | 0.103 | -0.022 | 0.229 |
| Fl. plant comp. diss. ~ LAI diss. | 0.172 . | 0.046 | 0.299 |
| Fl. plant comp. diss. ~ fire seasonality (2-S) diss. | 0.255 * | 0.129 | 0.382 |
| Fl. plant comp. diss. ~ soil moisture diss. | 0.145 | 0.016 | 0.275 |
| Fl. plant comp. diss. ~ geographic dist. | 0.412 *** | 0.301 | 0.523 |
| LAI diss. ~ soil moisture diss. | 0.183 | 0.040 | 0.327 |
| LAI diss. ~ terrain diss. | 0.162 | 0.020 | 0.304 |
| LAI diss. ~ weather diss. | -0.156 | -0.299 | -0.013 |
| fire seas. (2-S) diss. ~ number fires diss. | 0.505 ** | 0.395 | 0.614 |
| fire seas. (2-S) diss. ~ soil moisture diss. | -0.210 * | -0.334 | -0.086 |
| number fires diss. ~ land context diss. | 0.385 . | 0.253 | 0.517 |
| number fires diss. ~ terrain diss. | 0.052 | -0.089 | 0.193 |
| number fires diss. ~ geographic dist. | 0.091 | -0.048 | 0.230 |
| soil moisture diss. ~ weather diss. | 0.145 | -0.001 | 0.291 |
| soil moisture diss. ~ geographic dist. | 0.026 | 0.122 | 0.174 |
| land context diss. ~ terrain diss. | -0.180 | -0.324 | -0.035 |
| land context diss. ~ geographic dist. | -0.101 | -0.247 | 0.045 |
| terrain diss. ~ geographic dist. | 0.085 | -0.064 | 0.233 |

**β-diversity due to species turnover ($\beta_{ST}$)**

Table S.5.2. Coefficients path analysis for interaction β-diversity due to species turnover ($\beta_{ST}$). We used permutation-based path analysis (Fourtune et al. 2018). Levels of significance are noted as: . < 0.1, * <0.05, ** < 0.01, ***<0.001.

| Path | Std. Estimate | Lower CI | Upper CI |
|---|---|---|---|
| $\beta_{ST}$ ~ flowering plant abundance dissimilarity | 0.475 *** | 0.363 | 0.587 |
| $\beta_{ST}$ ~ flowering plant composition dissimilarity | 0.172 * | 0.044 | 0.300 |
| $\beta_{ST}$ ~ vegetation structure dissimilarity | -0.042 | -0.149 | 0.064 |
| $\beta_{ST}$ ~ LAI diss. | 0.147 | 0.039 | 0.255 |
| $\beta_{ST}$ ~ fire seasonality (2-S) diss | -0.079 | -0.205 | 0.047 |
| $\beta_{ST}$ ~ number fires dissimilarity | 0.194 . | 0.064 | 0.323 |
| $\beta_{ST}$ ~ land context diss. | 0.074 | -0.043 | 0.191 |
| $\beta_{ST}$ ~ geographic distance | 0.126 . | 0.006 | 0.247 |
| Fl. plant abund. diss. ~ Fl. plant comp. diss. | 0.286 ** | 0.142 | 0.430 |
| Fl. plant abund. diss. ~ fire seasonality (2-S) diss. | 0.105 | -0.033 | 0.244 |
| Fl. plant abund. diss. ~ soil moisture diss. | 0.009 | -0.126 | 0.145 |
| Fl. plant abund. diss. ~ land context diss. | 0.201 . | 0.071 | 0.331 |
| Fl. plant abund. diss. ~ weather diss. | 0.188 * | 0.061 | 0.315 |
| Fl. plant abund. diss. ~ geographic dist. | 0.201 * | 0.061 | 0.340 |
| Fl. plant comp. diss. ~ veg. structure diss. | 0.103 | -0.022 | 0.229 |
| Fl. plant comp. diss. ~ LAI diss. | 0.172 | 0.046 | 0.299 |
| Fl. plant comp. diss. ~ fire seasonality (2-S) diss. | 0.255 * | 0.129 | 0.382 |
| Fl. plant comp. diss. ~ soil moisture diss. | 0.145 | 0.016 | 0.275 |
| Fl. plant comp. diss. ~ geographic dist. | 0.412 *** | 0.301 | 0.523 |
| LAI diss. ~ soil moisture diss. | 0.183 | 0.040 | 0.327 |
| LAI diss. ~ terrain diss. | 0.162 | 0.020 | 0.304 |
| LAI diss. ~ weather diss. | -0.156 | -0.299 | -0.013 |
| fire seas. (2-S) diss. ~ number fires diss. | 0.505 ** | 0.395 | 0.614 |
| fire seas. (2-S) diss. ~ soil moisture diss. | -0.210 * | -0.334 | -0.086 |
| number fires diss. ~ land context diss. | 0.385 * | 0.253 | 0.517 |
| number fires diss. ~ terrain diss. | 0.052 | -0.089 | 0.193 |
| number fires diss. ~ geographic dist. | 0.091 | -0.048 | 0.230 |
| soil moisture diss. ~ weather diss. | 0.145 | -0.001 | 0.291 |
| soil moisture diss. ~ geographic dist. | 0.026 | 0.122 | 0.174 |
| land context diss. ~ terrain diss. | -0.180 | -0.324 | -0.035 |
| land context diss. ~ geographic dist. | -0.101 | -0.247 | 0.045 |
| terrain diss. ~ geographic dist. | 0.085 | -0.064 | 0.233 |

**Rewiring ($\beta_{RW}$)**

Table S.5.3. Coefficients path analysis for rewiring ($\beta_{RW}$). We used permutation-based path analysis (Fourtune et al. 2018). Levels of significance are noted as: . < 0.1, * <0.05, ** < 0.01, ***<0.001.

| Path | Std. Estimate | Lower CI | Upper CI |
|---|---|---|---|
| $\beta_{RW}$ ~ flowering plant abundance dissimilarity | -0.447 *** | -0.567 | -0.328 |
| $\beta_{RW}$ ~ vegetation structure dissimilarity | 0.018 | -0.099 | 0.136 |
| $\beta_{RW}$ ~ LAI diss. | -0.186 . | -0.303 | -0.069 |
| $\beta_{RW}$ ~ fire seasonality (2-S) diss | 0.084 | -0.054 | 0.222 |
| $\beta_{RW}$ ~ number fires dissimilarity | -0.222 . | -0.365 | -0.078 |
| $\beta_{RW}$ ~ land context diss. | -0.052 | -0.182 | 0.079 |
| $\beta_{RW}$ ~ geographic distance | -0.156 * | -0.281 | -0.032 |
| Fl. plant abund. diss. ~ Fl. plant comp. diss. | 0.286 *** | 0.142 | 0.430 |
| Fl. plant abund. diss. ~ fire seasonality (2-S) diss. | 0.105 | -0.033 | 0.244 |
| Fl. plant abund. diss. ~ soil moisture diss. | 0.009 | -0.126 | 0.145 |
| Fl. plant abund. diss. ~ land context diss. | 0.201 . | 0.071 | 0.331 |
| Fl. plant abund. diss. ~ weather diss. | 0.188 . | 0.061 | 0.315 |
| Fl. plant abund. diss. ~ geographic dist. | 0.201 * | 0.061 | 0.340 |
| Fl. plant comp. diss. ~ veg. structure diss. | 0.103 | -0.022 | 0.229 |
| Fl. plant comp. diss. ~ LAI diss. | 0.172 | 0.046 | 0.299 |
| Fl. plant comp. diss. ~ fire seasonality (2-S) diss. | 0.255 * | 0.129 | 0.382 |
| Fl. plant comp. diss. ~ soil moisture diss. | 0.145 | 0.016 | 0.275 |
| Fl. plant comp. diss. ~ geographic dist. | 0.412 *** | 0.301 | 0.523 |
| LAI diss. ~ soil moisture diss. | 0.183 | 0.040 | 0.327 |
| LAI diss. ~ terrain diss. | 0.162 | 0.020 | 0.304 |
| LAI diss. ~ weather diss. | -0.156 | -0.299 | -0.013 |
| fire seas. (2-S) diss. ~ number fires diss. | 0.505 ** | 0.395 | 0.614 |
| fire seas. (2-S) diss. ~ soil moisture diss. | -0.210 * | -0.334 | -0.086 |
| number fires diss. ~ land context diss. | 0.385 * | 0.253 | 0.517 |
| number fires diss. ~ terrain diss. | 0.052 | -0.089 | 0.193 |
| number fires diss. ~ geographic dist. | 0.091 | -0.048 | 0.230 |
| soil moisture diss. ~ weather diss. | 0.145 | -0.001 | 0.291 |
| soil moisture diss. ~ geographic dist. | 0.026 | 0.122 | 0.174 |
| land context diss. ~ terrain diss. | -0.180 | -0.324 | -0.035 |
| land context diss. ~ geographic dist. | -0.101 | -0.247 | 0.045 |
| terrain diss. ~ geographic dist. | 0.085 | -0.064 | 0.233 |

### MONTHLY TRENDS OF SPATIAL Β-DIVERSITY

Table S.5.4. Best-fit models for multiple linear regressions of monthly metrics of interaction β-diversity and its partition within preserves with sampling coverage > 0.6 against monthly flowering plant characteristics (same models as original). Levels of significance are noted as: . < 0.1, * <0.05, ** < 0.01, ***<0.001. We excluded an outlier in the model for $\beta_{RW}$.

| | **Fl. Pl. Abundance** | **Chao 1 Diversity** | **Fl. Pl. Richness** | **Fl. Pl. Evenness** | **Fl. Pl. Abundance x Chao 1 Diversity** | **Fl. Pl. Abundance x Fl. Pl. Evenness** | **Fl. Pl. Richness x Fl. Pl. Evenness** | **Adj. $R^2$** |
|---|---|---|---|---|---|---|---|---|
| $\beta_{Int}$ | -0.040 * | - | 0.019 . | -0.015 | - | -0.031 * | 0.004 | 0.040 |
| $\beta_{ST}$ | -0.077 ** | - | 0.042 | - | - | - | - | 0.093 |
| $\beta_{RW}$ | 0.075 *** | 0.013 | - | - | -0.062 ** | - | - | 0.217 |

**Supplementary material 6. Permutation-based path analysis models for presence/absence data**

**PATH ANALYSIS: BIOGEOGRAPHICAL AND ECOLOGICAL DRIVERS**

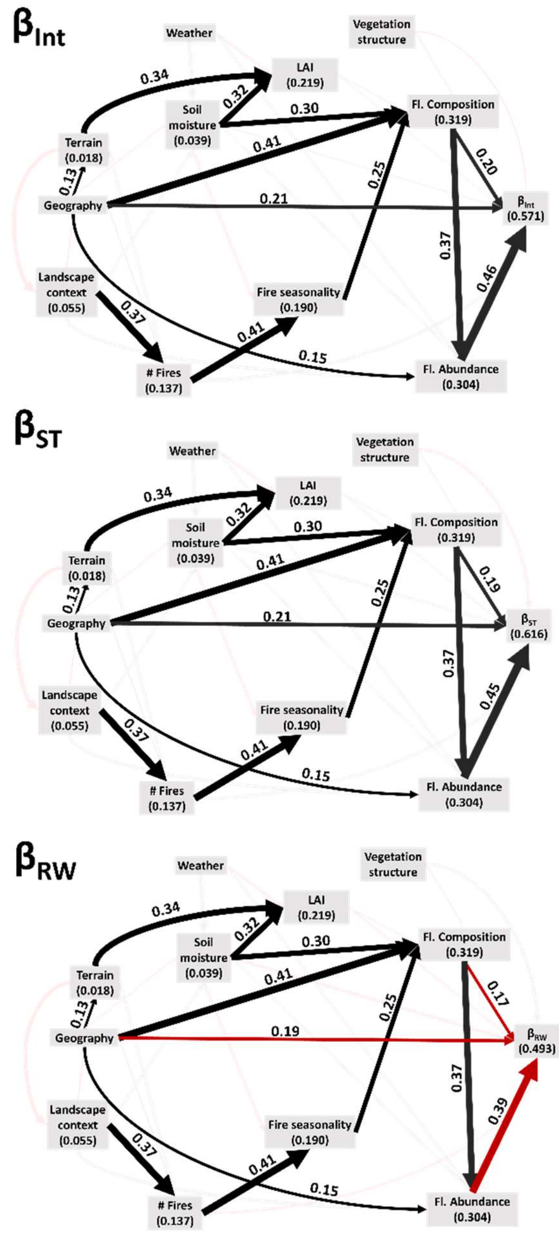

Figure S.6. 1. Path analysis for $\beta_{Int}$, $\beta_{ST}$, and $\beta_{RW}$ based on presence/absence data. The names in the boxes refer to site dissimilarities in respect to that particular factor. Positive relationships are portrayed by black lines and indicate parallel trends of both connecting variables (e.g., two sites with similar landscape context have similar number of fires). Negative relationships are shown in red and indicate opposite trends of both connecting variables (e.g., sites with similar soil moisture have different fire seasonality). Bold lines represent significant effects while faded lines represent not significant ones.

### Plant-pollinator interactions ($\beta_{Int}$)

Table S.6.1. Coefficients path analysis for plant-pollinator interactions ($\beta_{Int}$). We used permutation-based path analysis (Fourtune et al. 2018). Levels of significance are noted as: . < 0.1, * <0.05, ** < 0.01, ***<0.001.

| Path | Std. Estimate | Lower CI | Upper CI |
|---|---|---|---|
| $\beta_{Int}$ ~ flowering plant abundance dissimilarity | 0.464 *** | 0.379 | 0.549 |
| $\beta_{Int}$ ~ flowering plant composition dissimilarity | 0.201 ** | 0.106 | 0.297 |
| $\beta_{Int}$ ~ vegetation structure dissimilarity | -0.060 | -0.137 | 0.018 |
| $\beta_{Int}$ ~ LAI diss. | 0.092 | 0.010 | 0.173 |
| $\beta_{Int}$ ~ number fires dissimilarity | 0.152 . | 0.074 | 0.230 |
| $\beta_{Int}$ ~ soil moisture diss. | 0.063 | -0.023 | 0.149 |
| $\beta_{Int}$ ~ geographic distance | 0.205 ** | 0.119 | 0.290 |
| Fl. plant abund. diss. ~ Fl. plant comp. diss. | 0.368 *** | 0.261 | 0.476 |
| Fl. plant abund. diss. ~ soil moisture diss. | 0.147 . | 0.042 | 0.252 |
| Fl. plant abund. diss. ~ land context diss. | 0.167 . | 0.069 | 0.265 |
| Fl. plant abund. diss. ~ weather diss. | 0.135 . | 0.036 | 0.234 |
| Fl. plant abund. diss. ~ geographic dist. | 0.151 * | 0.042 | 0.260 |
| Fl. plant comp. diss. ~ veg. structure diss. | 0.087 | -0.010 | 0.184 |
| Fl. plant comp. diss. ~ LAI diss. | 0.105 | 0.004 | 0.207 |
| Fl. plant comp. diss. ~ fire seasonality (2-S) diss. | 0.253 * | 0.157 | 0.350 |
| Fl. plant comp. diss. ~ soil moisture diss. | 0.296 ** | 0.196 | 0.397 |
| Fl. plant comp. diss. ~ geographic dist. | 0.410 *** | 0.323 | 0.496 |
| LAI diss. ~ soil moisture diss. | 0.316 * | 0.214 | 0.417 |
| LAI diss. ~ terrain diss. | 0.344 * | 0.245 | 0.442 |
| LAI diss. ~ weather diss. | -0.140 | -0.244 | -0.035 |
| fire seas. (2-S) diss. ~ number fires diss. | 0.410 ** | 0.312 | 0.507 |
| fire seas. (2-S) diss. ~ soil moisture diss. | -0.149 | -0.254 | -0.044 |
| number fires diss. ~ land context diss. | 0.373 ** | 0.268 | 0.478 |
| number fires diss. ~ terrain diss. | 0.143 | 0.031 | 0.255 |
| number fires diss. ~ geographic dist. | 0.064 | -0.047 | 0.174 |
| soil moisture diss. ~ weather diss. | 0.182 | 0.069 | 0.295 |
| soil moisture diss. ~ geographic dist. | -0.090 . | -0.205 | 0.025 |
| land context diss. ~ terrain diss. | -0.216 . | -0.329 | -0.103 |
| land context diss. ~ geographic dist. | -0.068 | -0.183 | 0.048 |
| terrain diss. ~ geographic dist. | 0.133 * | 0.018 | 0.248 |

**$\beta$-diversity due to species turnover ($\beta_{ST}$)**

Table S.6.2. Coefficients path analysis for interaction $\beta$-diversity due to species turnover ($\beta_{ST}$). We used permutation-based path analysis (Fourtune et al. 2018). Levels of significance are noted as: . < 0.1, * <0.05, ** < 0.01, ***<0.001.

| Path | Std. Estimate | Lower CI | Upper CI |
|---|---|---|---|
| $\beta_{ST}$ ~ flowering plant abundance dissimilarity | 0.454 *** | 0.379 | 0.549 |
| $\beta_{ST}$ ~ flowering plant composition dissimilarity | 0.194 * | 0.106 | 0.297 |
| $\beta_{ST}$ ~ vegetation structure dissimilarity | -0.113 . | -0.206 | -0.060 |
| $\beta_{ST}$ ~ LAI diss. | 0.124 | 0.047 | 0.201 |
| $\beta_{ST}$ ~ number fires dissimilarity | 0.136 | 0.061 | 0.210 |
| $\beta_{ST}$ ~ soil moisture diss. | 0.136 | 0.054 | 0.217 |
| $\beta_{ST}$ ~ geographic distance | 0.214 *** | 0.132 | 0.295 |
| Fl. plant abund. diss. ~ Fl. plant comp. diss. | 0.368 *** | 0.261 | 0.476 |
| Fl. plant abund. diss. ~ soil moisture diss. | 0.147 . | 0.042 | 0.252 |
| Fl. plant abund. diss. ~ land context diss. | 0.167 . | 0.069 | 0.265 |
| Fl. plant abund. diss. ~ weather diss. | 0.135 | 0.036 | 0.234 |
| Fl. plant abund. diss. ~ geographic dist. | 0.151 * | 0.042 | 0.260 |
| Fl. plant comp. diss. ~ veg. structure diss. | 0.087 | -0.010 | 0.184 |
| Fl. plant comp. diss. ~ LAI diss. | 0.105 | 0.004 | 0.207 |
| Fl. plant comp. diss. ~ fire seasonality (2-S) diss. | 0.253 * | 0.157 | 0.350 |
| Fl. plant comp. diss. ~ soil moisture diss. | 0.296 * | 0.196 | 0.397 |
| Fl. plant comp. diss. ~ geographic dist. | 0.410 *** | 0.323 | 0.496 |
| LAI diss. ~ soil moisture diss. | 0.316 * | 0.214 | 0.417 |
| LAI diss. ~ terrain diss. | 0.344 * | 0.245 | 0.442 |
| LAI diss. ~ weather diss. | -0.140 | -0.244 | -0.035 |
| fire seas. (2-S) diss. ~ number fires diss. | 0.410 ** | 0.312 | 0.507 |
| fire seas. (2-S) diss. ~ soil moisture diss. | -0.149 | -0.254 | -0.044 |
| number fires diss. ~ land context diss. | 0.373 ** | 0.268 | 0.478 |
| number fires diss. ~ terrain diss. | 0.143 | 0.031 | 0.255 |
| number fires diss. ~ geographic dist. | 0.064 | -0.047 | 0.174 |
| soil moisture diss. ~ weather diss. | 0.182 | 0.069 | 0.295 |
| soil moisture diss. ~ geographic dist. | -0.090 . | -0.205 | 0.025 |
| land context diss. ~ terrain diss. | -0.216 . | -0.329 | -0.103 |
| land context diss. ~ geographic dist. | -0.068 | -0.183 | 0.048 |
| terrain diss. ~ geographic dist. | 0.133 * | 0.018 | 0.248 |

**Rewiring ($\beta_{RW}$)**

Table S.6.3. Coefficients path analysis for rewiring ($\beta_{RW}$). We used permutation-based path analysis (Fourtune et al. 2018). Levels of significance are noted as: . < 0.1, * <0.05, ** < 0.01, ***<0.001.

| Path | Std. Estimate | Lower CI | Upper CI |
|---|---|---|---|
| $B_{RW}$ ~ flowering plant abundance dissimilarity | -0.391 *** | -0.485 | -0.297 |
| $B_{RW}$ ~ flowering plant composition dissimilarity | -0.165 * | -0.269 | -0.061 |
| $B_{RW}$ ~ vegetation structure dissimilarity | 0.151 . | 0.067 | 0.234 |
| $B_{RW}$ ~ LAI diss. | -0.123 | -0.211 | -0.035 |
| $B_{RW}$ ~ number fires dissimilarity | -0.110 | -0.195 | -0.025 |
| $B_{RW}$ ~ soil moisture diss. | -0.152 | -0.245 | -0.059 |
| $B_{RW}$ ~ geographic distance | -0.190 ** | -0.283 | -0.097 |
| Fl. plant abund. diss. ~ Fl. plant comp. diss. | 0.368 *** | 0.261 | 0.476 |
| Fl. plant abund. diss. ~ soil moisture diss. | 0.147 . | 0.042 | 0.252 |
| Fl. plant abund. diss. ~ land context diss. | 0.167 . | 0.069 | 0.265 |
| Fl. plant abund. diss. ~ weather diss. | 0.135 . | 0.036 | 0.234 |
| Fl. plant abund. diss. ~ geographic dist. | 0.151 * | 0.042 | 0.260 |
| Fl. plant comp. diss. ~ veg. structure diss. | 0.087 | -0.010 | 0.184 |
| Fl. plant comp. diss. ~ LAI diss. | 0.105 | 0.004 | 0.207 |
| Fl. plant comp. diss. ~ fire seasonality (2-S) diss. | 0.253 * | 0.157 | 0.350 |
| Fl. plant comp. diss. ~ soil moisture diss. | 0.296 ** | 0.196 | 0.397 |
| Fl. plant comp. diss. ~ geographic dist. | 0.410 *** | 0.323 | 0.496 |
| LAI diss. ~ soil moisture diss. | 0.316 * | 0.214 | 0.417 |
| LAI diss. ~ terrain diss. | 0.344 ** | 0.245 | 0.442 |
| LAI diss. ~ weather diss. | -0.140 | -0.244 | -0.035 |
| fire seas. (2-S) diss. ~ number fires diss. | 0.410 ** | 0.312 | 0.507 |
| fire seas. (2-S) diss. ~ soil moisture diss. | -0.149 | -0.254 | -0.044 |
| number fires diss. ~ land context diss. | 0.373 ** | 0.268 | 0.478 |
| number fires diss. ~ terrain diss. | 0.143 | 0.031 | 0.255 |
| number fires diss. ~ geographic dist. | 0.064 | -0.047 | 0.174 |
| soil moisture diss. ~ weather diss. | 0.182 | 0.069 | 0.295 |
| soil moisture diss. ~ geographic dist. | -0.090 . | -0.205 | 0.025 |
| land context diss. ~ terrain diss. | -0.216 . | -0.329 | -0.103 |
| land context diss. ~ geographic dist. | -0.068 | -0.183 | 0.048 |
| terrain diss. ~ geographic dist. | 0.133 * | 0.018 | 0.248 |

### MONTHLY TRENDS OF SPATIAL Β-DIVERSITY

Table S.6.4. Best-fit models for multiple linear regressions of monthly metrics of presence/absence interaction β-diversity and its partition within preserves against monthly flowering plant characteristics (same models as weighted). Levels of significance are noted as: . < 0.1, * <0.05, ** < 0.01, ***<0.001.

| | Fl. Pl. Abundance | Chao 1 Diversity | Fl. Pl. Richness | Fl. Pl. Evenness | Fl. Pl. Abundance x Chao 1 Diversity | Fl. Pl. Abundance x Fl. Pl. Evenness | Fl. Pl. Richness x Fl. Pl. Evenness | Adj. $R^2$ |
|---|---|---|---|---|---|---|---|---|
| $\beta_{Int}$ | -0.044 *** | - | 0.018 * | -0.013 | - | -0.032 ** | 0.003 | 0.062 |
| $\beta_{ST}$ | -0.043 ** | - | 0.018 | - | - | - | - | 0.120 |
| $\beta_{RW}$ | 0.031 *** | 0.005 | - | - | -0.019 * | - | - | 0.208 |

**Supplementary material 7. Boxplot of interaction beta diversity**

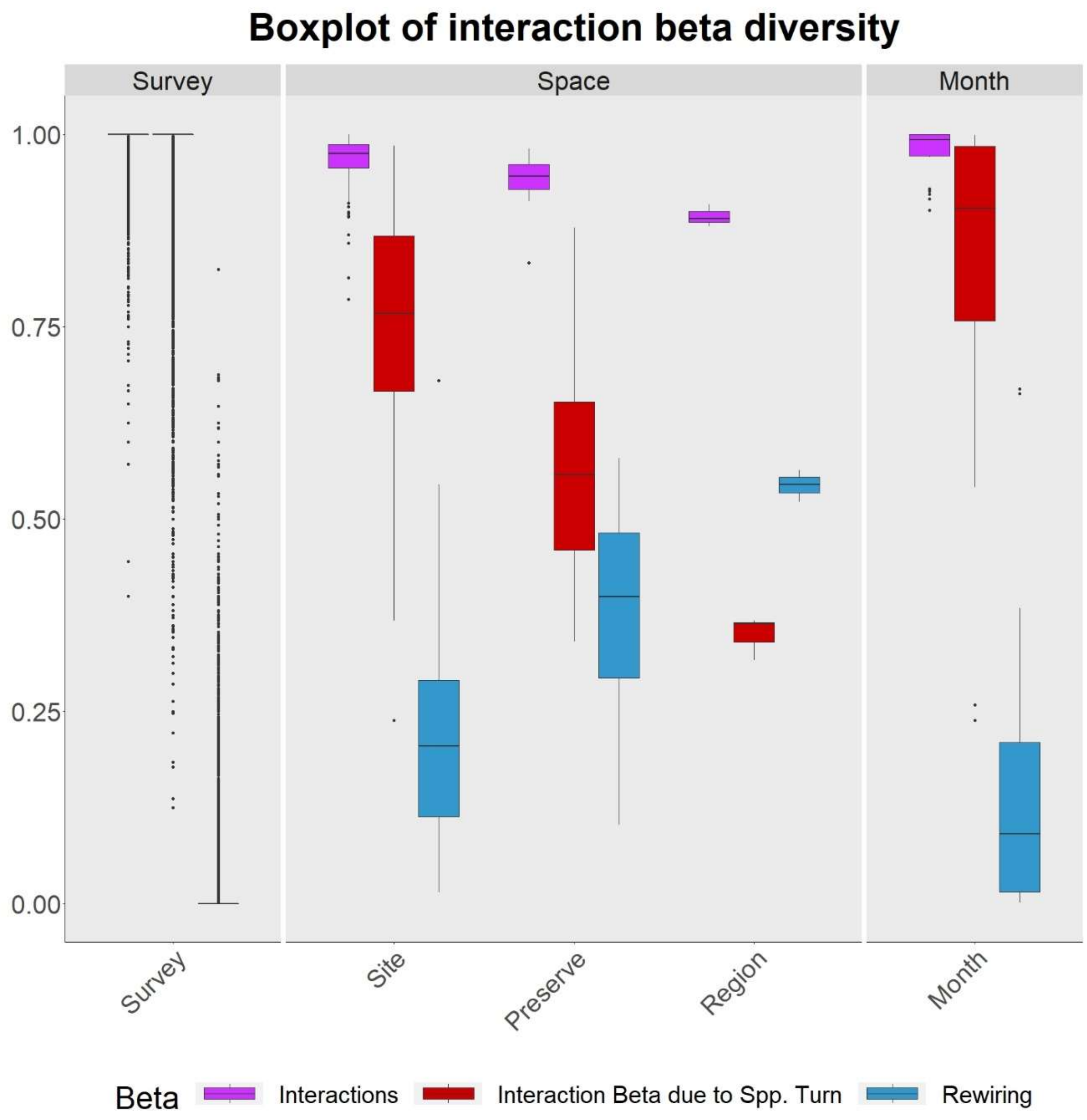

Figure S.7.1. Interaction β-diversity and its partition across surveys, sites, preserves, regions, and months. The three facets divide the plot among spatio-temporal (survey), spatial, and temporal (month) scales. The colors differ among metrics of β-diversity, magenta boxes represent interaction β-diversity (βInt), red boxes represent interaction β-diversity due to species turnover (βST), and blue boxes rewiring (βRW).
